# Detergent exchange from lipid nanoparticles into detergent micelles unlocks a tool for biochemical and kinetic characterization of membrane proteins

**DOI:** 10.64898/2025.12.23.696072

**Authors:** Rebecca S. Koweek, Hayley L. Knox, Greg J. Dodge, Barbara Imperiali, Karen N. Allen

**Affiliations:** Department of Chemistry, Boston University, Boston, MA USA 02215; Department of Biology and Department of Chemistry, Massachusetts Institute of Technology, Cambridge, MA USA 02139

**Keywords:** Membrane protein, lipid nanoparticles, detergent micelle, amphiphilic copolymers, membrane protein purification strategy

## Abstract

Bacterial membrane proteins make up ∼30% of the prokaryotic genome and play key roles in infection and virulence. Membrane protein chemistry has advanced in recent years, including purification strategies that mimic “native-like” lipid environments, such as lipid nanoparticles, amphipols, and nanodiscs. The use of styrene maleic acid co-polymers to form a lipid nanoparticle has become increasingly common in membrane protein purification, especially for proteins which are not amenable to detergent extraction from the cellular membrane fraction. Yet, for some biochemical and biophysical methods it is preferable to use detergent-solubilized protein. Here we show a general exchange method to transfer membrane proteins from lipid nanoparticles to detergent micelles while retaining protein fold, homogeneity and function. Conditions were first optimized for co-polymer dispersion and recovery into detergents, and analytical methods employed to assess activity and quality of detergent-solubilized proteins. Twelve protein targets were purified in co-polymer based on a 16-polymer screen. This selection was followed by an eight-detergent screen in the presence of calcium ions for optimal dissolution of the nanoparticle, producing detergent-stabilized protein. In all membrane proteins assessed, homogeneity and folding were retained from the initial purification in lipid nanoparticles through the detergent-exchange protocol. For membrane enzymes that have proven to be experimentally intractable once detergent solubilized, we were able to observe catalytic activity using the detergent-exchanged material. The use of this protocol to purify membrane proteins provides great versatility for biochemical and kinetic characterization than was previously accessible.

## Introduction

In both prokaryotic and eukaryotic organisms, membrane proteins make up roughly 30% of all cellular proteins and are involved in essential biological processes, including signaling, transport, and the maintenance of cellular structural integrity.^1,2^ Additionally, an estimated 70% of all drugs target membrane proteins.^3^ Structurally, they can be classified as peripheral (membrane-associated) or integral (embedded within the cell membrane). Integral membrane proteins are especially valuable targets, however, this protein class is challenging to purify *in vitro* and general strategies have proven elusive.

Biochemical studies of membrane proteins typically require solubilization from the cell membrane. Detergents are commonly used to extract and purify membrane proteins, but not all proteins are amendable to this strategy.^4^ Work on membrane proteins has advanced in recent years, including purification with molecules that mimic “native-like” lipid environments, such as lipid nanoparticles, amphipols, and nanodiscs.^5–7^ Previous work by Dodge *et. al* tested styrene maleic acid co-polymers (SMALPs) to extract proteins encompassing a variety of membrane topologies, that were recalcitrant to extraction using detergent micelles.^8,9^ In collaboration with Cube Biotech GmbH, an extensive screening protocol was developed for assessing purity and quality of membrane protein purified in lipid nanoparticles.^10^ With this methodology, a platform for generating selective nanobodies for a panel of membrane proteins was implemented.^11^ A recent review of membrane proteins purified in SMALPs describes multiple proteins that are unstable when purified in detergents but stable when purified in lipid nanoparticles.^12^ The use of SMALPs and other lipid nanoparticles has become increasingly common in membrane-protein purification and this method has been successful for structural analysis via cryogenic electron microscopy (cryo-EM).^9,13,14^

To expand the toolkit for studying membrane proteins that cannot be extracted directly into detergent micelles, we sought to develop a general exchange method to transfer membrane proteins from lipid nanoparticles to detergent micelles. This strategy can be used for proteins that do not extract directly from the cell-envelope fraction into detergent micelles but where further biochemical experiments in detergent are desired. The resulting protein preparation provides additional options for kinetic optimization and analysis, including biochemical assays and steady-state kinetic analysis. The method also enables X-ray crystallographic studies as proteins purified in SMALPs are not compatible with traditional crystallographic methods.

To verify the generalizability of this method, a panel of proteins were selected, representing various protein folds and topologies of membrane association including the monotopic membrane enzymes *Helicobacter pullorum* PglC and *Neisseria gonorrhoeae* PglB; the membrane-associated enzyme *Campylobacter concisus* PglI, and the transmembrane proteins *Salmonella enterica* Wzx, *E. coli* WcaJ, *S. enterica* WbaP, *Streptococcus pneumoniae* CpsE, *Simonsiella muelleri* BiPGT1, *Aeromonas hydrophila* WecP and *Myxococcales bacterium* GT1 (**Fig. 1**).^15,16^ The proteins are grouped as follows: monotopic, membrane-associated, and transmembrane. The enzymes chosen are involved in bacterial glycoconjugate biosynthesis and selected from bacterial pathogens such as *E. coli, S. enterica*, and *N. gonorrhoeae*. Glycoconjugates play a crucial role in bacterial cell-wall stability and in mediating pathogen-host interactions, rendering them prime targets for inhibiton.^17^ These enzymes are difficult to purify and characterize *in vitro*, highlighting a need for a strategy to generate high quality, stable protein. Initially in this protocol, a sixteen-polymer screen is utilized to identify the optimal co-polymer for homogeneity and yield of each membrane protein.^10^ A larger-scale SMALP purification is subsequently completed to enable an eight-detergent exchange screen (n-dodecyl-β-D-maltoside (DDM), N,N-dimethyldodecylamine N-oxide (LDAO), 3-[(3-cholamidopropyl)dimethylammonio]-1-propanesulfonate (CHAPS), n-decyl-β-D-maltoside (DM), n-dodecyl-phosphocholine (Fos-12), n-tetradecyl-N,N-dimethyl-3-ammonio-1-propanesulfonate (ANZERGENT 3-14), 5-cyclohexyl-1-pentyl-β-D-maltoside (CYMAL-5), and n-octyl-β-D-glucopyranoside (OG)), where the lipid nanoparticle is first destabilized by high salt and the membrane protein is recovered in detergent micelles (**Fig. 2**).

**Figure 1.**
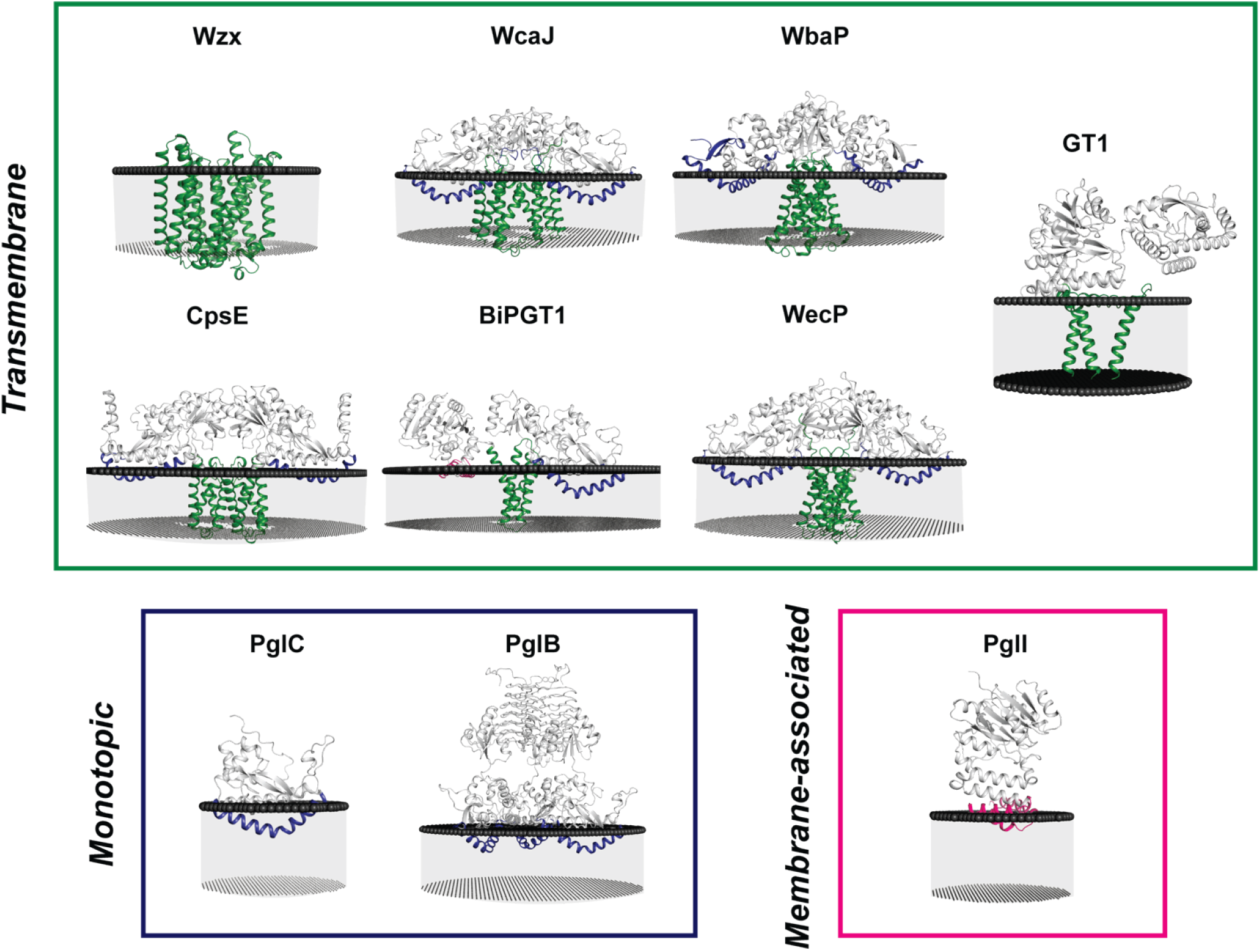
Target proteins for detergent exchange protocol. Prokaryotic membrane proteins, models generated in AlphaFold3 (except for WbaP, where PDB ID: 8TB3 was used) and modeled into membranes (black spheres) using the PPM3 server. Proteins are colored based on membrane interaction (blue: reentrant membrane helix, green: transmembrane, pink: membrane association, and grey: no interaction). Proteins are grouped based on membrane protein topology: transmembrane (*S. enterica* Wzx (P26400), *E. coli* WcaJ (P71241), *S. enterica* WbaP (P26406), *S. pneumoniae* CpsE (Q8KWP9), *S. muelleri* BiPGT1 (V9HM87), *A. hydrophila* WecP (B3FN88), *Myxococcales bacterium* GT1 (A0A7Y3BPP4); green frame), monotopic (*H. pullorum* PglC (E1B268), *N. gonorrhoeae* PglB (Q5FAE1); blue frame), membrane-associated (*C. concisus* strain 13826 PglI (A7ZEU0); pink frame). BiPGT2 could not be docked in the PPM server as the interdomain orientation is not known (**Fig. S1**).

**Figure 2.**
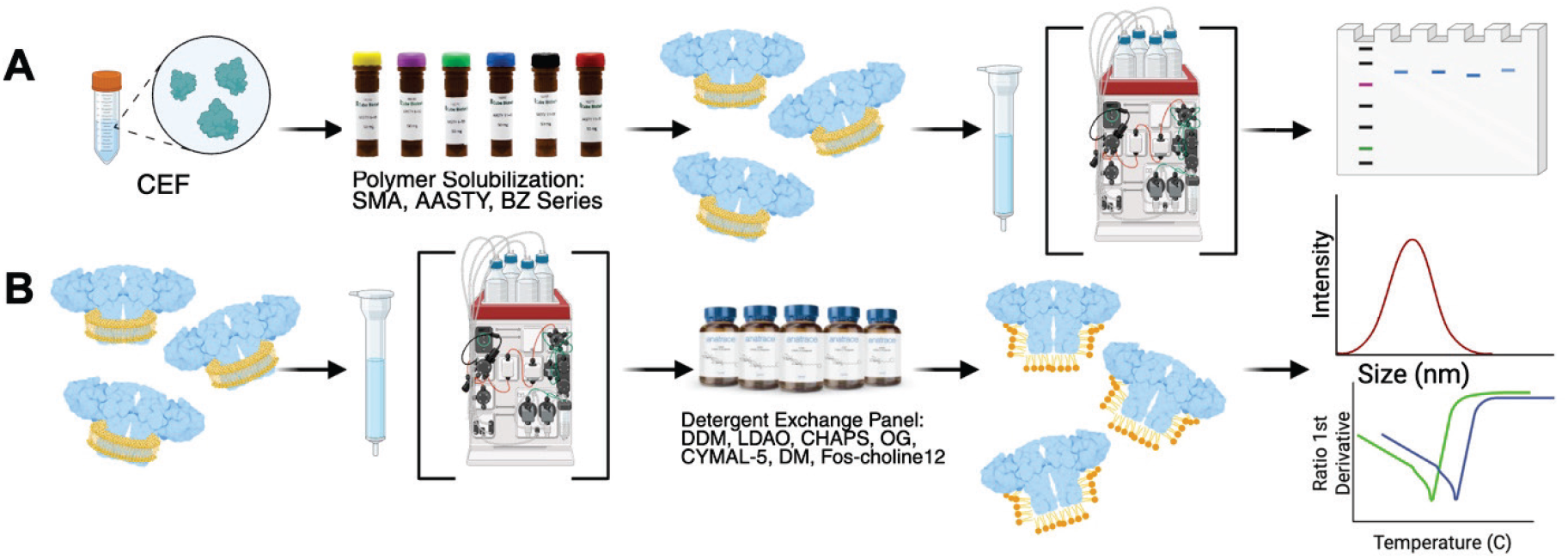
Detergent exchange protocol for membrane proteins. (A) Membrane protein purification into SMA or AASTY co-polymers. Concentrated cell-envelope fraction (CEF) is combined with co-polymer to extract membrane proteins. The proteins are purified over a Streptactin-XT column and can be further purified using size-exclusion chromatography. Protein in lipid nanoparticle is analyzed using SDS-PAGE, dynamic light scattering, and unfolding analysis. (B) Membrane proteins solubilized in lipid nanoparticle are purified over a Streptactin-XT column, further purified using size-exclusion chromatography, then exchanged into detergent micelles resulting in pure detergent-solubilized protein analyzed as in (A).

This strategy is resource and time efficient and is applicable to membrane proteins of various topologies and cell-membrane association modalities, with consistently high-quality proteins obtained through the purification and exchange process. Five proteins, representing each membrane topology category, were shown to be fully active following detergent exchange. For two enzymes within the study, PglC from *H. pullorum* and PglI from *C. concisus*, extraction and purification in DDM micelles was compared to purification in lipid nanoparticles followed by detergent-exchange into DDM micelles. In both cases, the detergent-exchanged protein was of higher homogeneity than that extracted directly into detergent, and protein quality remained the same. This work establishes the detergent exchange method as a reliable and consistent method for high-quality protein production suitable for downstream biochemical analysis.

## Methods

### Protein Expression and Purification

Plasmids were constructed with tags optimized in a previously described purification strategy.^8^ In short, plasmids include a SUMO tag to aid with solubility and a high-affinity dual-strep tag for purification. All proteins were full-length, with the exception of the construct for A0A315CH80 (BiPGT2), which encodes amino acids 138-965, omitting a disordered N-terminal region. Plasmids were transformed into C43 competent cells harboring pAM174, which encodes an arabinose-inducible Ulp1 protease for SUMO tag cleavage *in vivo*. Target proteins were expressed using auto-induction media^18^ in 0.5 L cultures, supplemented with 150 µg/mL of kanamycin (GoldBio) and 25 µg/mL of chloramphenicol (GoldBio) at 37 °C. At OD_600_ ∼0.6-0.8, 1 g of arabinose (GoldBio) was added to each flask and the temperature was lowered to 18 °C. Cells were harvested no later than 24 hours post-inoculation and pellets were stored at -80 °C.

For cell membrane preparation, cell pellets were resuspended at ∼3 mL/g with Buffer A (50 mM HEPES pH 7.5, 300 mM NaCl), 40 mg of lysozyme (GoldBio), 10 mg of DNase (GoldBio), and 50 µL of protease inhibitor cocktail (Millipore Sigma, Abcam). The mixture was resuspended at room temperature for 45 minutes. All subsequent purification steps were performed at 4 °C unless otherwise stated. Cells were lysed with a microfluidizer using 1 pass at 18,000 psi. Lysate was clarified by centrifugation in a Ti70 rotor at 9,300 RPM for 45 minutes (or 8,000 RPM in a Ti45 rotor) to remove cell debris. The resulting supernatant was centrifuged in a Ti70 rotor at 36,100 RPM for 65 minutes (or 33,000 RPM in a Ti45 rotor) to separate the membrane and soluble fraction. Supernatant was discarded and membrane pellets were resuspended in Buffer A using a glass dounce homogenizer to shear the cell membranes. Resuspended membrane cell pellets are referred to as cell envelope fraction (CEF). Total protein concentration in CEF was measured using absorbance at 280 nm. CEF was diluted to 50 mg/mL, flash frozen in 5 mL aliquots in LN_2_, and stored at -80 °C.

For purification of proteins in copolymer for subsequent exchange into detergents, one aliquot of 5 mL CEF was thawed from -80 °C at room temperature, then solubilized with 4 mL of Buffer A and 1 mL of 10% co-polymer (Cube Biotech) for 1 hour at room temperature. The reaction was centrifuged for 65 minutes at 42,000 RPM in a Ti70 rotor. A gravity column of Streptactin-XT resin (IBA) with a 1 mL bed volume was equilibrated with 15 mL of Buffer A, and the supernatant was run over the column bed twice at room temperature. The column was washed with 10-15 mL of Buffer A followed by 3 mL of Buffer B which was incubated on the resin bed for 10-30 minutes at room temperature before eluting. The eluted protein was concentrated to ∼500 µL with an Amicon filter (Millipore Sigma) concentrator (MW cutoff 10k Da or 30k Da, depending on the protein) and stored on ice at 4 °C. Protein concentration ranged between 1 and 5 mg/mL.

Larger-scale purification of detergent-exchanged membrane proteins was required for kinetic analysis. CEF (10 mL at 50 mg/mL) combined with 1.1 mL of 10% co-polymer (Cube Biotech) for extraction was solubilized at room temperature for 1 hour. Solubilized CEF was centrifuged for 65 minutes at 42,000 RPM in a Ti70 rotor, and the supernatant was run over the column bed twice over a 1 mL Streptactin-XT (IBA) column equilibrated in Buffer A at room temperature. The column was washed with 10-15 mL of Buffer A and incubated for >30 min with 4 mL of Buffer B before eluting. The eluent was concentrated to <500 µL and injected onto a Superose 6 (Cytiva) sizing column equilibrated with Buffer C (25 mM HEPES pH 7.5, 150 mM NaCl). Peak fractions were combined and exchanged into the previously determined optimal detergent, typically 2.5 mL of purified protein, 312.5 µL of 10% detergent (Anatrace, GoldBio), and 312.5 µL of 200 CaCl_2_ + 200 mM MgCl_2_ [detergent exchange reaction occurs with final detergent concentration at 1%, to ensure all detergents exceed CMC]. The reaction mix was rotated at 4 °C for 1 hour, then centrifuged for 45 min at 42,000 RPM in a Ti70 rotor. The resultant purified protein exchanged into detergent was concentrated to <2.5 mL and desalted using an equilibrated PD-10 column (Cytiva). Desalting buffer varied based on the detergent, but generally was composed of 25 mM HEPES pH 7.5, 150 mM NaCl, and 5% glycerol, with a detergent concentration above the CMC. Desalted protein was concentrated to <300 µL and flash frozen in LN_2_.

### Polymer screen

Membrane proteins were screened with a sixteen-co-polymer screen using the method described in the Cube Biotech technical note.^10^ Briefly, 75 µL of CEF was solubilized with 10% solutions of each co-polymer and solubilized at room temperature for 1 hour. Solubilized CEF was centrifuged in a TLA-100 rotor at 35,000 RPM for 35 minutes, then incubated with 40 µL of equilibrated Streptactin-XT resin for 30 minutes. Streptactin-XT beads were washed 3 times with 600 µL of Buffer A, centrifuging in a tabletop centrifuge (Eppendorf 5415D) for 30 s at 700 x g to pellet the beads after each wash. Buffer B (50 mM HEPES pH 7.5, 300 mM NaCl, 50 mM biotin) was added (30 µL) and incubated at room temperature for >10 min. Modified SDS-PAGE analysis (with denaturation at 45 °C for 30 minutes^19^) was performed on each sample to assess purity, as well as measurement of the radius of gyration and thermal unfolding using a Prometheus PANTA (NanoTemper).

### Detergent exchange screen

Exchange of proteins from lipid nanoparticles into detergent micelles was performed in 50 µL reactions. For eight detergents, purified membrane proteins in lipid nanoparticles ranging from 1 to 5 mg/mL were combined with 10% (w/v) detergent and a high salt solution (10 mM CaCl_2_ + 10 mM MgCl_2_). The mixture was incubated at 4 °C for 1 hr, then centrifuged at 35,000 RPM for 35 min in a TLA-100 rotor to separate co-polymer from detergent-solubilized protein. Modified SDS-PAGE analysis was performed on each sample (with denaturation at 45 °C for 30 minutes^19^), as well as radius of gyration and thermal unfolding using a Prometheus PANTA (NanoTemper).

### Protein Thermostability and Activity Assays

Thermostability and particle size were assessed using a Prometheus PANTA (NanoTemper) for protein samples in lipid nanoparticle and in detergent micelles. Thermostability was determined by monitoring 330 and 350 nm, with a temperature increase of 1.5 °C/second.

Promega UMP-Glo (Cat #VA1130) assays for *S. enterica* WbaP, *T. thermophilus* WbaP, *E. coli* WcaJ and *H. pullorum* PglC were performed using established procedures.^20^ Briefly, enzyme at 1 µM was added to a mixture of UndP at 20 µM and assay buffer (50 mM HEPES pH 7.5, 100 mM NaCl, 5 mM MgCl_2_, 0.1% TritionX-100) and initiated with 1 µL of substrate (UDP-galactose for *S. enterica* WbaP and *T. thermophilus* WbaP, UDP-glucose for *E. coli* WcaJ, and UDP-N,N’-diacetylbacillosamine (UDP-diNAcBac) for *H. pullorum* PglC) at a final concentration of 20 µM. For *S. enterica* WbaP and *E. coli* WcaJ, reactions were performed with final soluble substrate concentrations of 20, 50, and 100 µM. Reactions were performed in an 11 µL reaction volume, run at ambient temperature for 30 min and quenched directly with a 1:1 ratio of UMP-Glo reagent in a 96-well plate (Corning) for luminescence measurement on a Synergy H1 plate reader (Agilent Technologies). Luminescence measurements were converted to [UMP] with a standard curve.

For *C. jejuni* PglI, the glycosylation activity was monitored by the UDP-Glo assay from Promega (Cat #V6963) as previously described.^21^ Briefly, 2.8 µM PglI, 1 mM UDP-Glc, assay buffer (0.1% Triton, 50 mM HEPES pH 7.5, 100 mM NaCl, 5 mM MgCl_2_), and acceptor (UndP-Bac-(GalNAc)_5_) at an unknown concentration was added. The acceptor was chemoenzymatically synthesized as detailed in Lukose *et al*.^22^ PglI and approximately 70 µM UndP-Bac-(GalNAc)_5_ were incubated for 10 minutes prior to the addition of the sugar donor UDP-Glc. The PglI reaction was performed in an 11 µL volume and quenched into 11 µL of UDP detection reagent after 30 min. The samples were transferred to a 96 well plate (Corning), shaken at low speed for 30 s and incubated for 1 hr at 25 °C prior to measuring the luminescence on a Synergy H1 plate reader (Agilent Technologies).

## Results

To develop a generalizable method to transfer membrane-embedded enzymes from co-polymer nanoparticles to detergent, we first optimized conditions for co-polymer dispersion and recovery into detergents, then used analytical methods to assess activity and quality of detergent-solubilized product. This was achieved for proteins of various membrane topologies (**Fig. 1**). Co-polymers vary based on the ratio of acid to styrene or acrylic acid unit, and detailed analysis has shown that the resulting particles differ in their stability to divalent cations.^23–25^ Conditions were empirically determined for two selected proteins to ensure that the lipid nanoparticles were destabilized prior to transfer to detergent micelles. Purified proteins were incubated with varying concentrations (5-10 mM) of MgCl_2_, CaCl_2_, or both, analyzed by SDS-PAGE and the hydrodynamic radius determined to assess lipid nanoparticle destabilization. For both proteins purified in SMA200 or AASTY 11-50, a solution of 10 mM MgCl_2_ and 10 mM CaCl_2_ was the most effective at destabilizing the proteins (**Fig. S2**). Following detergent exchange, excess Ca^2+^ and Mg^2+^ salts were removed via a desalting column.

Successful exchange of proteins from lipid nanoparticle into detergent micelles was observed for all membrane protein topologies: monotopic (*H. pullorum* PglC, *N. gonorrhoeae* PglB), membrane-associated (*C. concisus* strain 13826 and *C. jejuni* PglI), and transmembrane (*E. coli* WcaJ, *S. enterica* WbaP, *T. thermophilus* WbaP, *S. pneumoniae* CpsE, *Aeromonas hydrophila* WecP, *Simonsiella muelleri* bifunctional monoPGT (BiPGT1), *Limnohabitans sp. Jir61* bifunctional monoPGT (BiPGT2), *Myxococcales bacterium* glycosyltransferase (GT1), and *S. enterica* Wzx (formerly RfbX)). Each target was purified in an optimal co-polymer determined by a 16-polymer screen using the dual Strep tag for high purity (**Figs. S3-S11**), then mixed with high salt and one of eight different detergents to allow dissolution of the nanoparticle and stabilization by detergent. In two cases (BiPGT1 and BiPGT2; **Figs. S6 and S7**), the low purification yield precluded reliable interpretation of the SDS-PAGE, thus DLS alone was used to select the best polymer. The detergents screened were selected to sample different chemical scaffolds and properties and represent two major detergent classes used for membrane protein solubilization: non-ionic (DDM, DM, ANZERGENT 3-14, OG, CYMAL-5) and zwitterionic (LDAO, CHAPS, Fos-12). Each detergent-exchanged reaction was centrifuged to yield the detergent-solubilized material in the supernatant (soluble fraction) which was analyzed using SDS-PAGE, DLS, and a thermal unfolding assay (**Figs. S12-S23**). In many cases the protein quality was inferior, which was only apparent by DLS which was used to select the best polymer and detergent. Ultimately, protein quality as evaluated by DLS confirms successful exchange into detergent micelles as assessed by a radius shift from ∼10 nm to ∼4 nm.

Target protein purification was successfully scaled up for biochemical studies (**Fig. 3**). Yields varied between proteins, but generally ∼ 30% of pure protein was successfully exchanged from the initial purification step into lipid nanoparticle and size-exclusion chromatography (**Table 1**). The resultant protein is highly homogenous and suitable for biochemical characterization. We observed melting temperatures (T_m_) for most proteins by analysis of thermal unfolding measured via the complementary biophysical techniques of turbidity, scattering, and hydrodynamic radius. Most proteins in this study have multiple domains, which made analysis of unfolding using only the 350/330 nm absorbance ratio unreliable. Thus, T_m_ was measured using either the 350nm/330 nm absorbance ratio, or turbidity measurement; the T_m_ values typically ranged from ∼40-60 °C (**Table 1**).

**Table 1.** Membrane protein yield from detergent exchange. Melting temperature (T_m_) is measured using either ^a^ratio 350/330 nm absorbance or ^b^turbidity.

| Membrane Protein | UniProt ID | Protein fold | Yield from 10 mL CEF polymer purification (mg) | Yield from 10 mL CEF detergent exchange (mg) | Expected yield from 1L cell growth (mg) | Optimal detergent for exchange | Melting Temperature ($T_m$, °C) |
| --- | --- | --- | --- | --- | --- | --- | --- |
| *E. coli* WcaJ | P71241 | Large monoPGT | 3.0 | 1.0 | 3.0 | DDM | 55.7<sup>b</sup> |
| *S. enterica* WbaP | P26406 | Large monoPGT | 3.0 | 1.0 | 3.0 | DDM | 36.5<sup>a</sup> |
| *S. pneumoniae* CpsE | Q8KWP9 | Large monoPGT | 1.5 | 0.8 | 2.3 | Fos-12 | 54.5<sup>b</sup> |
| *A. hydrophila* WecP | B3FN88 | Large monoPGT | 2.7 | 0.9 | 2.7 | DM | 47.1<sup>a</sup> |
| *T. Thermophilus* WbaP | A0A9N7K5G2 | Large monoPGT | 1.5 | 0.8 | 2.3 | DDM | 59.8<sup>b</sup> |
| *Limnohabitans* sp. Jir61 BiPGT2 | A0A315CH80 | Bifunctional monoPGT | 1.5 | 0.4 | 1.0 | Fos-12 | 60.2<sup>a</sup> |
| *S. muelleri* BiPGT1 | V9HM87 | Bifunctional monoPGT | 1.5 | 0.5 | 1.5 | DM | 46.4<sup>b</sup> |
| *H. pullorum* PglC | E1B268 | Small monoPGT | 1.8 | 0.4 | 1.2 | DDM | 60.5<sup>a</sup> |
| *S. enterica* RfbX | P26400 | Transmembrane | 1.5 | 0.3 | 0.8 | CYMAL-5 | 57.9<sup>b</sup> |
| *N. gonorrhoeae* PglB | Q5FAE1 | Bifunctional monoPGT | 4.5 | 1.8 | 5.3 | DDM | 53.9<sup>b</sup> |
| *C. concisus* strain 13826 PglI | A7ZEU0 | Glycosyltransferase Fold-A | 3.0 | 0.9 | 3.0 | DDM | 48.2<sup>b</sup> |
| *C. jejuni* PglI | Q0P9C6 | Glycosyltransferase Fold-A | 3.0 | 0.9 | 3.0 | DDM | N/A |
| *Myxococcales* bacterium GT1 | A0A7Y3BPP4 | Bifunctional monoPGT | 1.5 | 0.2 | 0.6 | DM | 50.1<sup>b</sup> |

**Figure 3.**
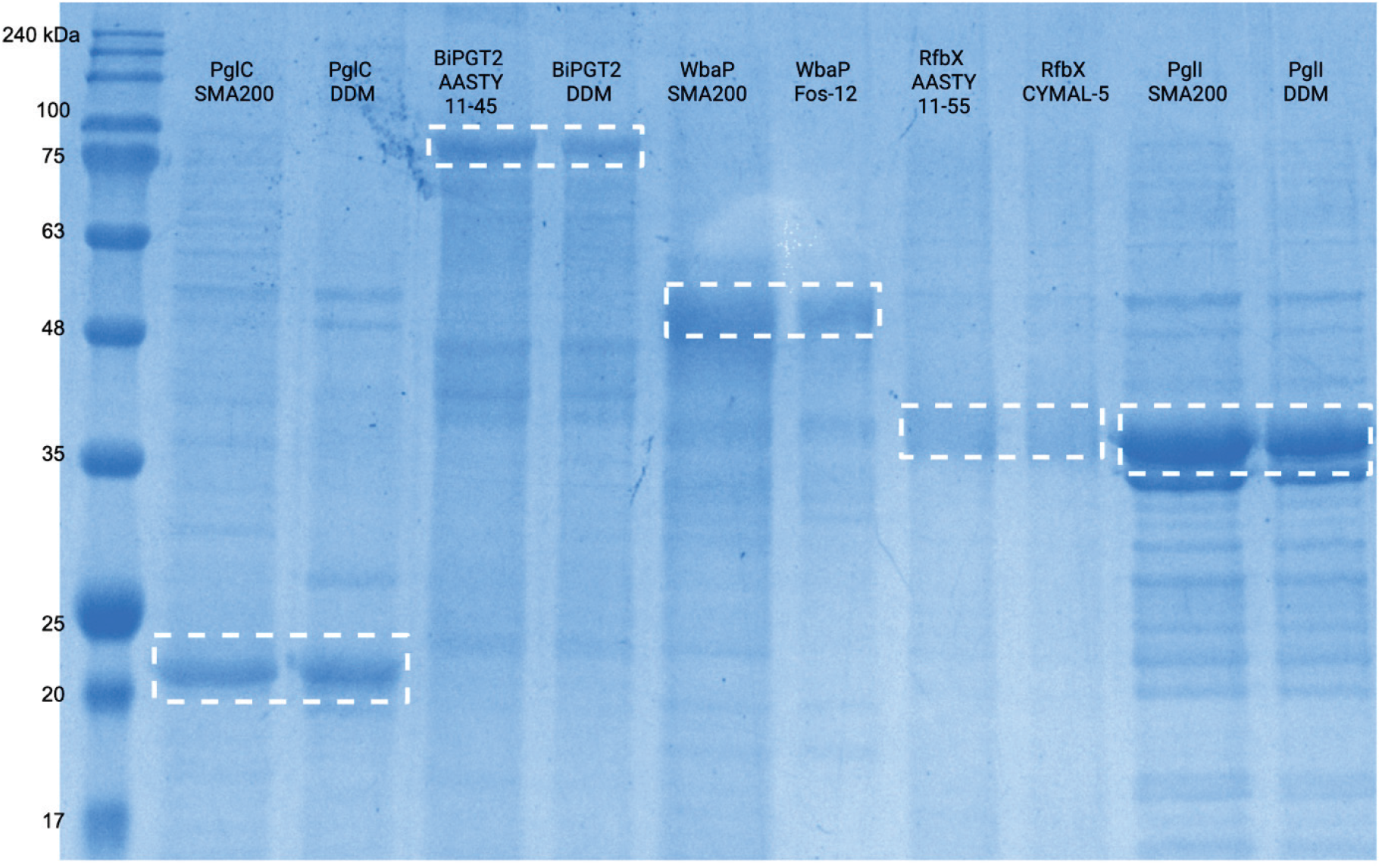
Representatives of four classes of membrane proteins purified in lipid nanoparticles and exchanged into detergent micelles. Membrane proteins from left to right with corresponding molecular weights: *H. pullorum* PglC in SMA200 and DDM (23.6 kDa), *Limnohabitans sp. Jir61* BiPGT2 in AASTY 11-45 and DDM (91.5 kDa), *S. enterica* WbaP in SMA200 and DDM (56.2 kDa), *S. enterica* RfbX in AASTY 11-55 and CYMAL-5 (48.6 kDa), and *C. jejuni* PglI in SMA200 and DDM (35.5 kDa). Membrane proteins with high pI values are known to run slightly below their expected molecular weight.

Purified proteins using the detergent-exchange method are comparable in quality to detergent-purified enzymes, as assessed by purity, T_m_, and homogeneity. Specific comparison was made for enzymes which could be purified directly into detergent but also purified in lipid nanoparticles: PglC from *H. pullorum* and PglI from *C. jejuni. H. pullorum* PglC could be purified in DDM detergent micelles and exchanged into DDM after purification in SMA200 (**Fig. 4A**). As compared to purification in detergent directly, when purified in lipid nanoparticle and exchanged into DDM micelles, the protein was of higher homogeneity, had the same T_m_, and retained enzymatic activity. PglI from *C. jejuni* was also purified successfully in DDM detergent micelles and can be exchanged into DDM after purification in SMA200 (**Fig. 4B**). The substrate of PglI from *C. jejuni* is known and the kinetics have been previously characterized.^22^ Notably, DLS data revealed that the protein purified in SMALP and then exchanged has higher homogeneity than when purified in DDM. Detergent-exchanged PglI displays a five-fold lower activity than the sample directly detergent purified. This is likely due to the use of CaCl_2_ for SMALPs dissolution, which may compete with the required Mg^2+^cofactor.

**Figure 4.**
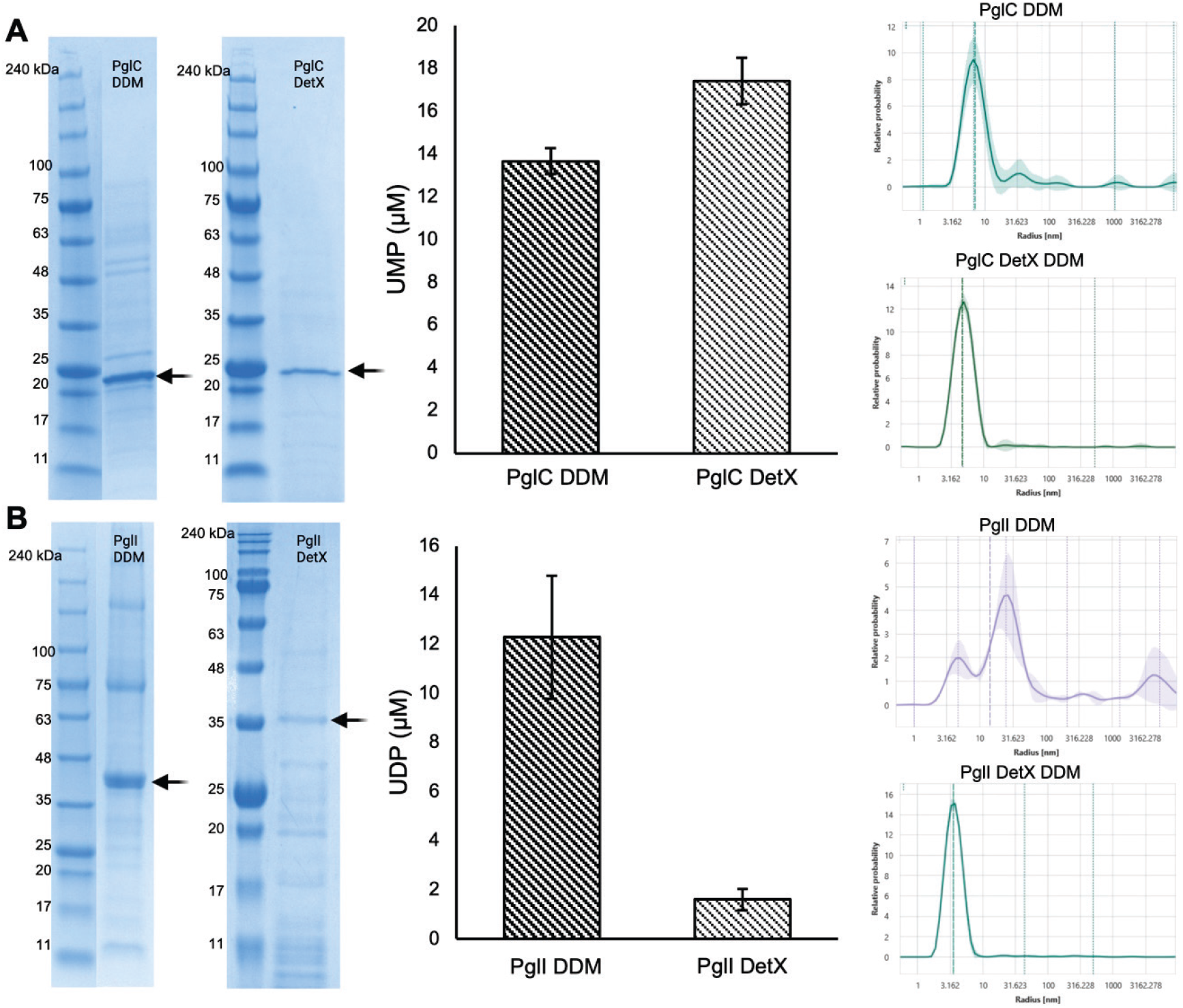
Comparison of detergent exchange with detergent purification. (**A**) **Left**: *H. pullorum* PglC purified in DDM micelles compared with PglC purified in SMALP 200 and exchanged into DDM (DetX). Arrow indicates target protein band. **Middle**: PglC endpoint activity using the UMP-Glo assay, showing that the enzyme does not lose activity as a result of the detergent exchange process. **Right**: DLS comparison between detergent-purified and detergent-exchanged PglC. (**B**) **Left**: *C. jejuni* PglI purified in DDM micelles compared with PglI purification in SMALP 200 and exchanged into DDM. Arrow indicates target protein band. **Middle**: PglI endpoint activity using the UDP-Glo assay. **Right**: DLS comparison between detergent-purified and detergent-exchanged PglI.

Enzymes of the large monoPGT subfamily of the monoPGT superfamily^26^ have been challenging to purify, making their *in vitro* kinetic analysis especially difficult. Previous studies demonstrated robust monoPGT activity and specificity in CEF when utilizing a radiolabeled assay.^8^ In this work, the UMP-Glo assay was used to confirm the *in vitro* activity of full-length, purified large monoPGTs *E. coli* WcaJ, *S. enterica* WbaP, and *T. thermophilus* WbaP. All three enzymes were purified in lipid nanoparticles and detergent-exchanged into DDM (**Fig. 5A**). Once exchanged into detergent, all three displayed activity with their cognate substrates (UDP-glucose for WcaJ, and UDP-galactose for both WbaP orthologs) as assessed via an endpoint assay (**Fig. 5B, C**). It has proven challenging to purify large monoPGTs in detergent which has served as a impediment in understanding their cellular activity. Importantly, these results demonstrate that the detergent exchange method may be used for determining specificity and kinetics of large monoPGTs that were previously inaccessible.

**Figure 5.**
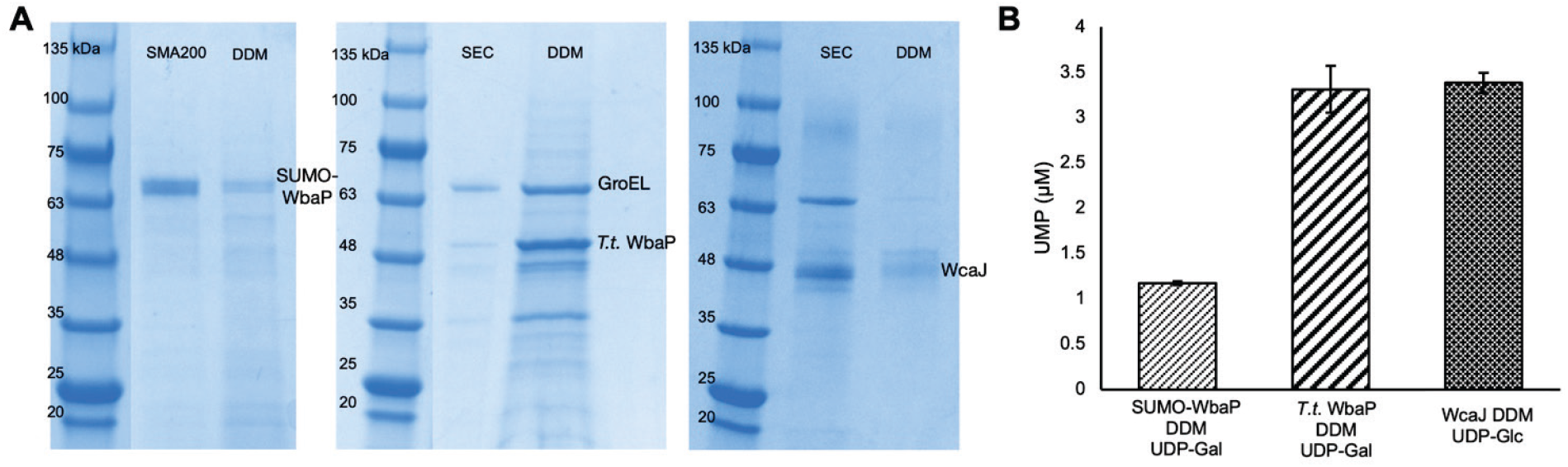
*In vitro* activity of *S. enterica* WbaP, *T. thermophilus* WbaP, and *E. coli* WcaJ. (**A**) SDS-PAGE of purification and exchange of WbaP orthologs and WcaJ into DDM micelles. (**B**) Activity of WbaP homologs in DDM with substrate UDP-galactose (UDP-Gal) and WcaJ in DDM with substrate UDP-glucose (UDP-Glc).

## Discussion

Our study has demonstrated that the detergent exchange method produces high-quality protein using a variety of analytical techniques. We assessed two proteins (PglC and PglI) that can be purified in detergent for direct comparison to detergent exchange and observed similar T_m_ values and higher homogeneity. For multiple proteins in the study (WcaJ and WbaP orthologs), we were able to assess activity *in vitro*, which requires production of folded and stable protein. For all membrane proteins assessed, we observed that homogeneity is retained from the initial purification into lipid nanoparticle through the detergent exchange protocol. We found that seven out of nine proteins in this study extracted in higher yield in SMA200 or the AASTY polymer series (specifically AASTY 11-50), whereas two proteins did not extract in high yield for any polymer (BiPGT1 and BiPGT2). As the proteins in this study span membrane topologies, the success with all proteins supports the generality of this method for applications which require homogeneous detergent-solubilized protein. Additionally, the detergent-exchange screen is an accessible tool for the identification of detergents that optimally stabilize membrane protein targets. In our hands, we observed than most proteins exchanged well into Fos-12, which has a structure similar to the phospholipids that comprise cell membranes, and DDM, which is commonly used for crystallographic structural studies.^27,28^

Membrane proteins that play critical roles in biological processes and are targets for development of therapeutic inhibitory ligands often require kinetic analysis. There has not yet been extensive kinetic characterization of membrane enzymes in lipid nanoparticles, compared to the widespread kinetic analysis of detergent-solubilized enzymes.^20^ Thus, for membrane enzymes that are experimentally intractable under traditional DDM conditions, detergent exchange offers an additional optimization strategy for biochemical assays of membrane-resident enzymes. This method provides an addition to multiple strategies for the biochemical, structural, and inhibitor analysis embodied by the extensive polymer screen and selective nanobody development previously established.^10,11^

Large monoPGT enzymes exemplify the utility of the detergent-exchange method, as they have proven difficult to purify via direct detergent extraction from CEF.^8^ This approach provides a direct method for kinetic analysis of large monoPGTs, which has been a roadblock to understanding their cellular action. Previous studies from the Valvano group showed that the catalytic domain of multiple large monoPGTs are active *in vivo*, and kinetic parameters were determined.^29,30^ Using detergent exchanged material herein we demonstrate enzymatic activity with three monoPGTs *in vitro* enabling future study of specificity and structure/function. The use of detergent-exchange to purify large monoPGTs provides great versatility for biochemical characterization than was previously accessible.

The study of membrane proteins has been hampered by the low yield upon heterologous expression and the loss of stability once extracted from the membrane. There have been significant recent advances in membrane protein solubilization, including the use of detergents, co-polymers, and amphipols.^31^ Co-polymers offer a versatile strategy for membrane protein purification by forming lipid nanoparticles around the protein. Such lipid nanoparticles are an excellent vehicle for membrane protein stabilization, purification, and manipulation^11^ but may have limitations in some structural and kinetic experiments. Lipid nanoparticle stability has been explored in terms of pH and cation sensitivity, but the destabilization of lipid nanoparticles is less understood. One previous study did succeed in destabilizing SMALPs and reconstituting the protein sample in detergent for mass spectrometry.^25^ However, lipid nanoparticles have since expanded to include both SMA and AASTY co-polymers, the latter of which form more robust lipid nanoparticles. This study expands on previous work to include an effective strategy for destabilizing and rescuing protein from the currently available co-polymers. The protocol incorporates extensive co-polymer and detergent screening to enable the study of recalcitrant membrane proteins and expand the current toolset for the study of targets of interest.

## Acknowledgements

This work was supported by NIH R01 GM039334 and R01 GM131627 (to K.N.A. and B.I.) and F32 GM149160 (to H.K.). We would like to thank Dr. Hugh Higinbotham for helpful discussions about activity assays.

## Notes

### Competing Interest Statement

The authors have declared no competing interest.

