## Supplementary material for "Detergent exchange from lipid nanoparticles into detergent micelles unlocks a tool for biochemical and kinetic characterization of membrane proteins": Table S1 and Figures S1-S23

Karen N. Allen – Department of Chemistry, Boston University, Boston, MA 02215, USA; Program in Biomolecular Pharmacology, Boston University School of Medicine, Boston, MA 02118, USA.

### Table of Contents


Figure S1. AlphaFold model for BiPGT2.

Figure S2. Destabilization of lipid nanoparticles for SMA and AASTY

Figure S3. Polymer screen data for *E. coli* WcaJ

Figure S4. Polymer screen data for *S. pneumoniae* CpsE

Figure S5. Polymer screen data for *S. enterica* RfbX

Figure S6. Polymer screen data for *Limnohabitans sp. Jir61* BiPGT2

Figure S7. Polymer screen data for *S. muelleri* BiPGT1

Figure S8. Polymer screen data for *C. concisus* PglI

Figure S9. Polymer screen data for *Myxococcales bacterium* GT1

Figure S10. Polymer screen data for *A. hydrophila* WecP

Figure S11. Polymer Screen for *T. thermophilus* WbaP

Figure S12. Detergent exchange screen data for *E. coli* WcaJ

Figure S13. Detergent exchange screen data for *S. pneumoniae* CpsE

Figure S14. Detergent exchange screen data for *S. enterica* WbaP

Figure S15. Detergent exchange screen data for *H. pullorum* PglC

Figure S16. Detergent exchange screen data for *S. enterica* RfbX

Figure S17. Detergent exchange screen data for *S. muelleri* BiPGT1

Figure S18. Detergent exchange screen data for *Limnohabitans sp. Jir61* BiPGT2

Figure S19. Detergent exchange screen data for *C. concisus* PglI

Figure S20. Detergent exchange screen data for *N. gonorrhoeae* PglB

Figure S21. Detergent exchange screen data for *Myxococcales bacterium* GT1

Figure S22. Detergent exchange screen data for *A. hydrophila* WecP

Figure S23. Detergent exchange screen data for *T. thermophilus* WbaP

Table S1. Primers for cloning plasmids

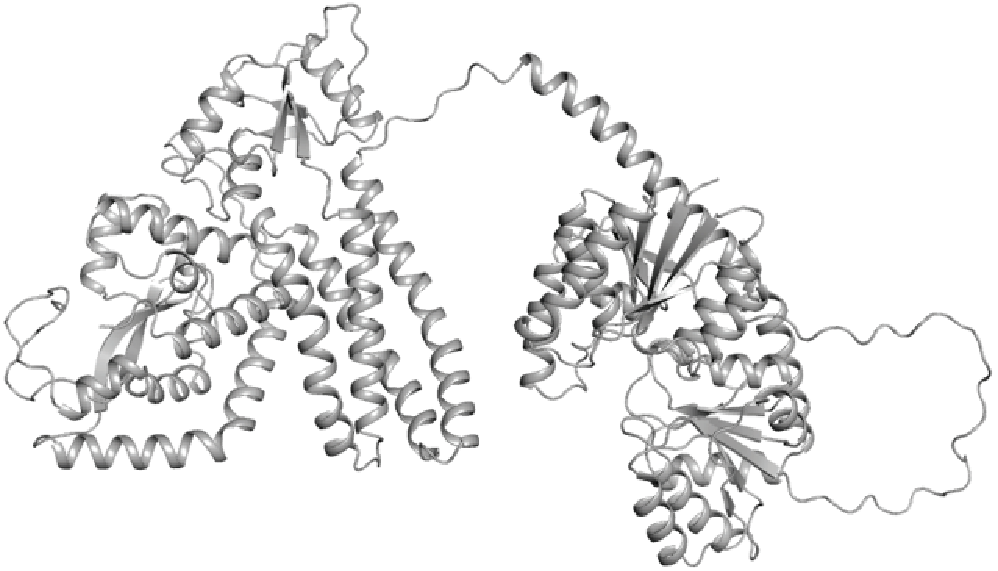

**Figure S1**. **AlphaFold model for BiPGT2.** UniProt ID: A0A315CH80. Although both the phosphoglycosyltransferase (left) and glycosyltransferase (right) domains are expected to be membrane bound, the interdomain relationship is unknown such that computational docking in the membrane leads to an unrealistic pose.

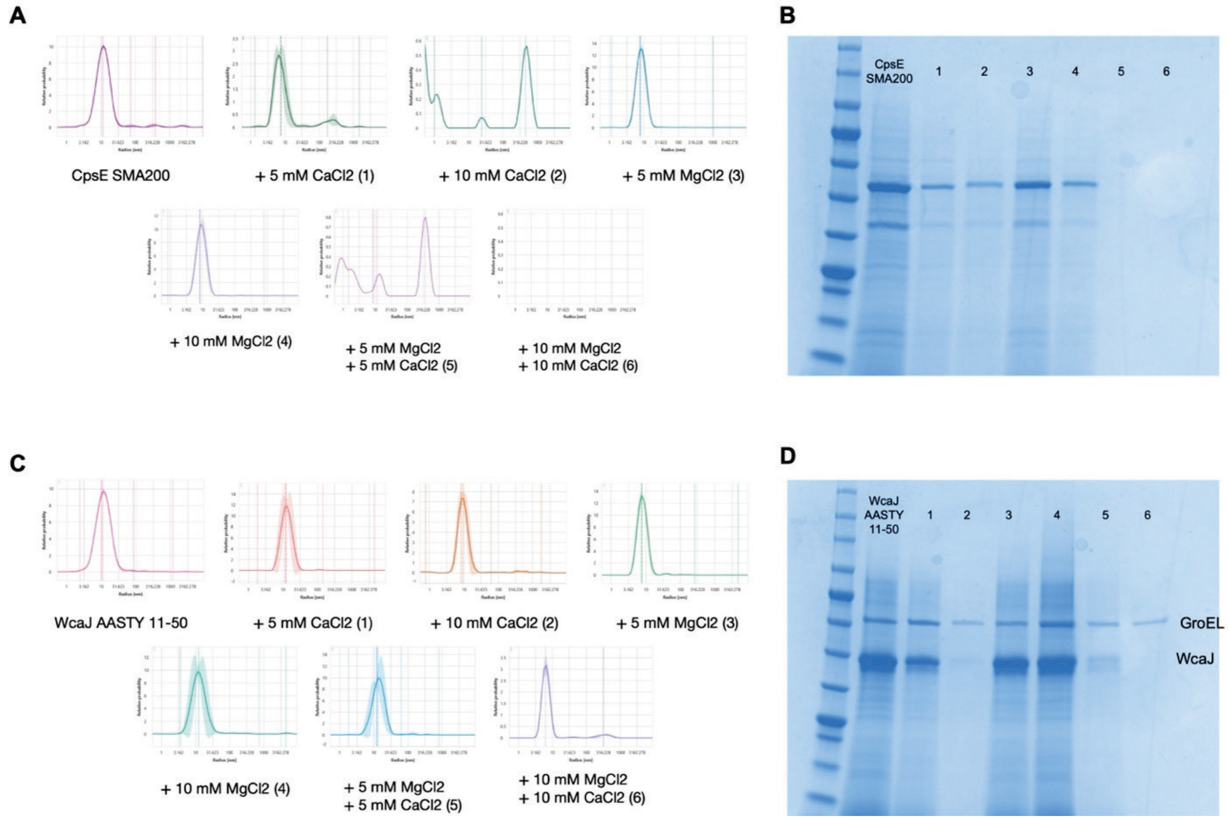


**Figure S2**. **Destabilization of lipid nanoparticles for SMA and AASTY**. **(A)** Radius shifts for CpsE in SMA200 show destabilization of lipid nanoparticle with varying salt concentrations. Lane numbers correspond to sample numbers **1**: 5 mM $CaCl_2$, **2**: 10 mM $CaCl_2$, **3**: 5 mM $MgCl_2$, **4**: 10 mM $MgCl_2$, **5**: 5 mM $CaCl_2$ + 5 mM $MgCl_2$, **6**: 10 mM $CaCl_2$ + 10 mM $MgCl_2$. **(B)** SDS-PAGE confirming loss of protein in lipid nanoparticle. **(C)** Radius shifts for WcaJ in AASTY 11-50 show destabilization of lipid nanoparticle with varying salt concentrations. **(D)** SDS-PAGE confirmed loss of protein in lipid nanoparticle. Lane numbers as in (**A**).

**A**

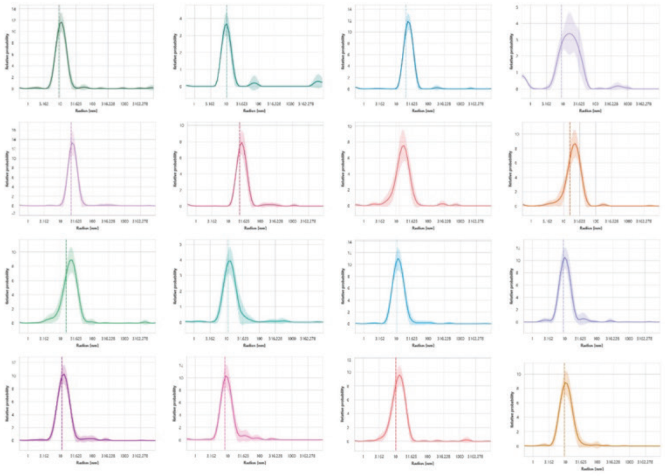

| Annotations | Cumulant Fit | | | |
|---|---|---|---|---|
| | Cumulant Radius (nm) | | Cumulant PDI | |
| Sample ID | ⌀ | σ | ⌀ | σ |
| Ec WcaJ SMA 25 | 9.84 | 0.34 | 0.26 | 0.05 |
| Ec WcaJ SMA 30 | 10.34 | 0.22 | 0.40 | 0.07 |
| Ec WcaJ SMA 40 | 20.98 | 0.44 | 0.29 | 0.03 |
| Ec WcaJ DIBMA | 9.02 | 0.74 | 0.83 | 0.05 |
| Ec WcaJ Sulfo-DIBMA | 22.24 | 0.50 | 0.22 | 0.04 |
| Ec WcaJ Sulfo-SMA | 23.25 | 0.59 | 0.30 | 0.04 |
| Ec WcaJ SMA BZ25 | 12.74 | 0.21 | 0.39 | 0.02 |
| Ec WcaJ SMA BZ30 | 15.89 | 0.43 | 0.40 | 0.03 |
| Ec WcaJ SMA BZ35 | 15.81 | 0.24 | 0.39 | 0.02 |
| Ec WcaJ SMA BZ40 | 11.37 | 0.25 | 0.32 | 0.04 |
| Ec WcaJ AASTY 6-45 | 10.71 | 0.08 | 0.22 | 0.02 |
| Ec WcaJ AASTY 11-45 | 9.32 | 0.08 | 0.25 | 0.02 |
| Ec WcaJ AASTY 6-50 | 11.39 | 0.12 | 0.25 | 0.01 |
| Ec WcaJ AASTY 11-50 | 8.39 | 0.07 | 0.30 | 0.02 |
| Ec WcaJ AASTY 6-55 | 10.46 | 0.34 | 0.32 | 0.05 |
| Ec WcaJ AASTY 11-55 | 9.50 | 0.08 | 0.25 | 0.02 |

**B**

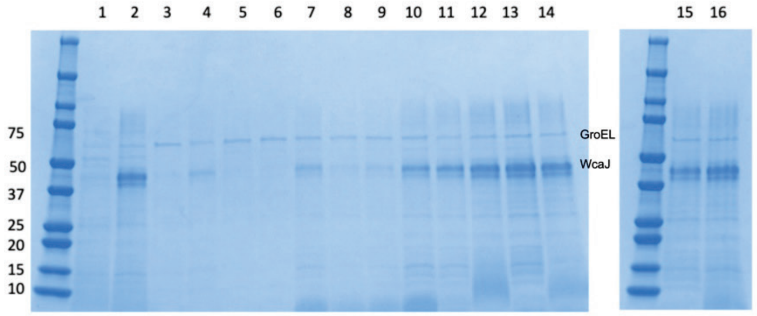


**Figure S3. Polymer screen data for *E. coli* WcaJ. (A)** Radius and polydispersity data for polymers in the following order: SMA100, SMA200, SMA300, DIBMA, Sulfo-DIBMA, Sulfo-SMA, SMA BZ25, SMA BZ30, SMA BZ35, SMA BZ40, AASTY 6-45, AASTY 11-45, AASTY 6-50, AASTY 11-50, AASTY 6-55, AASTY 11-55. **(B)** SDS-PAGE showing fractions from purification of WcaJ using the polymer screen. Lanes numbers correspond to samples: **1**: SMA100, **2**: SMA200, **3:** SMA300, **4**: DIBMA, **5**: Sulfo-DIBMA, **6**: Sulfo-SMA, **7**: SMA BZ25, **8**: SMA BZ30, **9**: SMA BZ35, **10**: SMA BZ40, **11:** AASTY 6-45, **12**: AASTY 11-45, **13**: AASTY 6-50, **14**: AASTY 11-50, **15**: AASTY 6-55, **16**: AASTY 11-55.

**A**

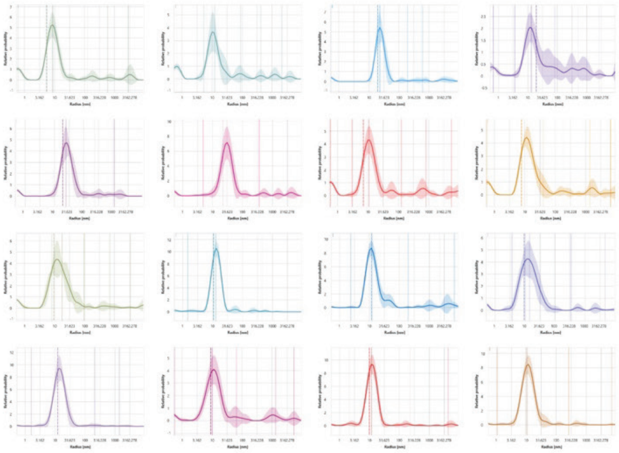

| Annotations | Cumulant Fit | | | |
|---|---|---|---|---|
| | Cumulant Radius (nm) | | Cumulant PDI | |
| Sample ID | ø | σ | ø | σ |
| CpsE SMA25 | 5.48 | 1.20 | 0.90 | 0.07 |
| CpsE SMA30 | 6.79 | 2.10 | 1.11 | 0.04 |
| CpsE SMA40 | 19.77 | 2.64 | 0.57 | 0.12 |
| CpsE DIBMA | 19.61 | 23.75 | 1.01 | 0.24 |
| CpsE Sulfo-DIBMA | 20.12 | 2.46 | 0.63 | 0.07 |
| CpsE Sulfo-SMA | 22.59 | 2.71 | 0.65 | 0.13 |
| CpsE SMALP BZ25 | 7.07 | 1.56 | 1.01 | 0.04 |
| CpsE SMALP BZ30 | 7.79 | 1.67 | 1.00 | 0.06 |
| CpsE SMALP BZ35 | 9.32 | 1.89 | 0.79 | 0.06 |
| CpsE SMALP BZ40 | 10.97 | 0.21 | 0.37 | 0.02 |
| CpsE AASTY 6-45 | 11.94 | 1.07 | 0.47 | 0.16 |
| CpsE AASTY 11-45 | 9.53 | 0.77 | 0.63 | 0.06 |
| CpsE AASTY 6-50 | 12.20 | 0.55 | 0.33 | 0.04 |
| CpsE AASTY 11-50 | 9.10 | 2.64 | 0.70 | 0.17 |
| CpsE AASTY 6-55 | 9.93 | 0.56 | 0.43 | 0.07 |
| CpsE AASTY 11-55 | 9.61 | 0.29 | 0.38 | 0.03 |

**B**

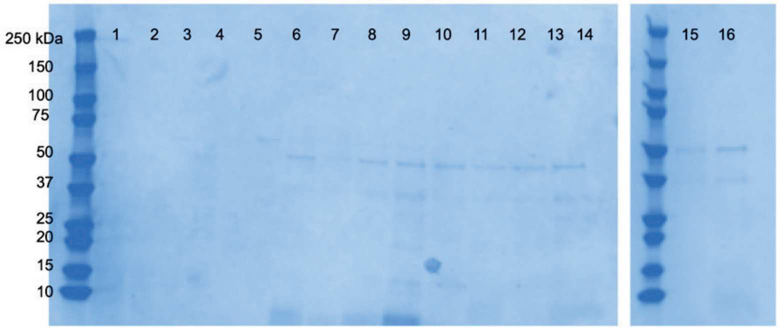


**Figure S4. Polymer screen data for *S. pneumoniae* CpsE. (A)** Radius and polydispersity data for polymers in the following order: SMA100, SMA200, SMA300, DIBMA, Sulfo-DIBMA, Sulfo-SMA, SMA BZ25, SMA BZ30, SMA BZ35, SMA BZ40, AASTY 6-45, AASTY 11-45, AASTY 6-50, AASTY 11-50, AASTY 6-55, AASTY 11-55. **(B)** SDS-PAGE of purification of CpsE using the polymer screen. Lane numbering is the same as Fig. S3.

**A**

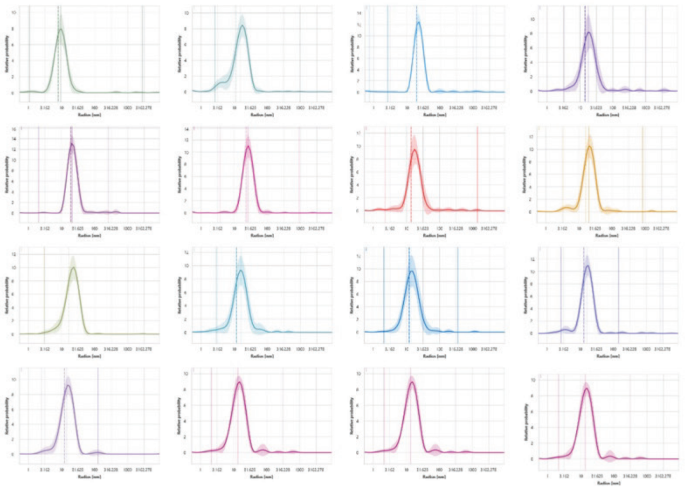

| Annotations | Cumulant Fit | | | |
|---|---|---|---|---|
| | Cumulant Radius (nm) | | Cumulant PDI | |
| Sample ID | ⌀ | σ | ⌀ | σ |
| RfbX SMA25 | 8.10 | 0.05 | 0.27 | 0.01 |
| RfbX SMA30 | 11.17 | 0.15 | 0.45 | 0.02 |
| RfbX SMA40 | 20.05 | 0.29 | 0.25 | 0.03 |
| RfbX DIBMA | 14.66 | 0.43 | 0.40 | 0.05 |
| RfbX Sulfo-DIBMA | 19.45 | 0.14 | 0.21 | 0.02 |
| RfbX Sulfo-SMA | 22.97 | 0.44 | 0.24 | 0.03 |
| RfbX SMALP BZ25 | 14.29 | 0.17 | 0.38 | 0.02 |
| RfbX SMALP BZ30 | 16.09 | 0.24 | 0.34 | 0.02 |
| RfbX SMALP BZ35 | 16.96 | 0.19 | 0.31 | 0.02 |
| RfbX SMALP BZ40 | 12.19 | 0.12 | 0.38 | 0.01 |
| RfbX AASTY 6-45 | 12.36 | 0.09 | 0.28 | 0.02 |
| RfbX AASTY 11-45 | 13.27 | 0.18 | 0.31 | 0.02 |
| RfbX AASTY 6-50 | 12.23 | 0.12 | 0.31 | 0.01 |
| RfbX AASTY 11-50 | 11.63 | 0.08 | 0.34 | 0.02 |
| RfbX AASTY 6-55 | 12.65 | 0.22 | 0.36 | 0.02 |
| RfbX AASTY 11-55 | 10.42 | 0.23 | 0.33 | 0.04 |

**B**

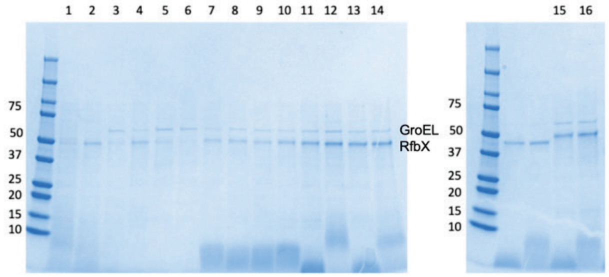


**Figure S5. Polymer screen data for *S. enterica* RfbX. (A)** Radius and polydispersity data for polymers in the following order: SMA100, SMA200, SMA300, DIBMA, Sulfo-DIBMA, Sulfo-SMA, SMA BZ25, SMA BZ30, SMA BZ35, SMA BZ40, AASTY 6-45, AASTY 11-45, AASTY 6-50, AASTY 11-50, AASTY 6-55, AASTY 11-55. **(B)** SDS-PAGE of purification of RfbX using the polymer screen. Lane numbering is the same as Fig. S3.

**A**

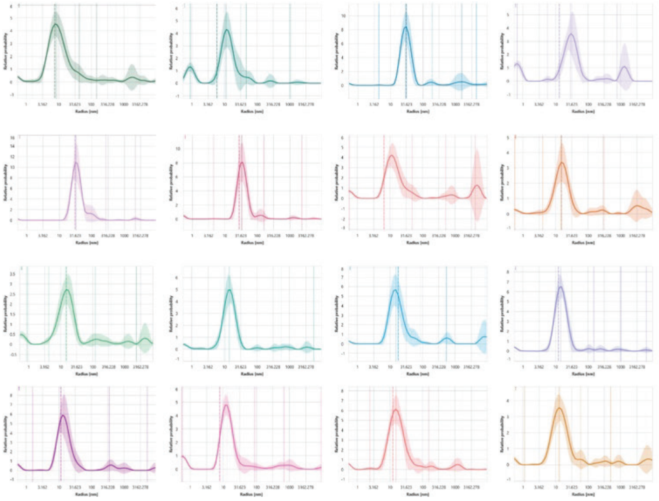

| Annotations | Cumulant Fit | | | |
|---|---|---|---|---|
| | Cumulant Radius (nm) | | Cumulant PDI | |
| H80 SMA25 | 7.56 | 1.41 | 0.68 | 0.12 |
| H80 SMA30 | 6.00 | 1.72 | 1.05 | 0.06 |
| H80 SMA40 | 30.98 | 12.26 | 0.48 | 0.29 |
| H80 DIBMA | 13.27 | 6.41 | 1.38 | 0.06 |
| H80 Sulfo-DIBMA | 31.03 | 1.05 | 0.34 | 0.06 |
| H80 Sulfo-SMA | 27.97 | 1.48 | 0.47 | 0.10 |
| H80 SMALP BZ25 | 6.56 | 3.76 | 0.59 | 0.34 |
| H80 SMALP BZ30 | 15.21 | 6.11 | 0.76 | 0.22 |
| H80 SMALP BZ35 | 16.38 | 13.03 | 1.01 | 0.19 |
| H80 SMALP BZ40 | 10.91 | 0.99 | 0.61 | 0.07 |
| H80 AASTY 6-45 | 17.89 | 9.97 | 0.61 | 0.31 |
| H80 AASTY 11-45 | 12.49 | 0.62 | 0.49 | 0.07 |
| H80 AASTY 6-50 | 11.41 | 2.10 | 0.71 | 0.11 |
| H80 AASTY 11-50 | 7.80 | 1.14 | 0.96 | 0.05 |
| H80 AASTY 6-55 | 12.48 | 0.77 | 0.62 | 0.06 |
| H80 AASTY 11-55 | 11.99 | 4.29 | 0.69 | 0.17 |

**Figure S6. Polymer screen data for *Limnohabitans sp. Jir61* BiPGT2. (A)** Radius and polydispersity data for polymers in the following order: SMA100, SMA200, SMA300, DIBMA, Sulfo-DIBMA, Sulfo-SMA, SMA BZ25, SMA BZ30, SMA BZ35, SMA BZ40, AASTY 6-45, AASTY 11-45, AASTY 6-50, AASTY 11-50, AASTY 6-55, AASTY 11-55. Due to low protein yield SDS-PAGE revealed faint bands precluding reliable interpretation, thus DLS data was used for designating the optimal polymer for BiPGT2.

**A**

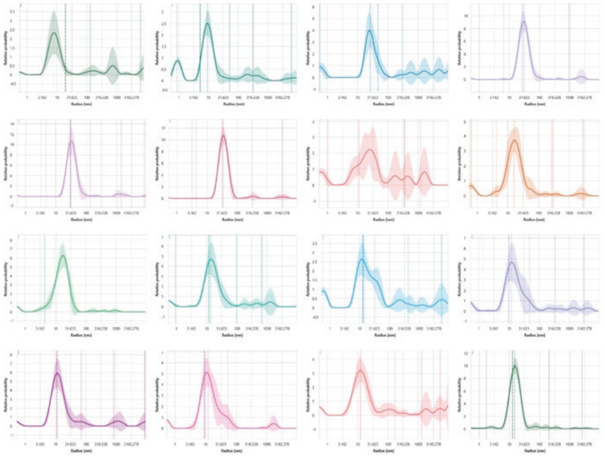

| Annotations | Cumulant Fit | | | |
|---|---|---|---|---|
| | Cumulant Radius (nm) | | Cumulant PDI | |
| Sample ID | ⌀ | σ | ⌀ | σ |
| M87 SMA 25 | 20.54 | 30.31 | 0.81 | 0.41 |
| M87 SMA 30 | 5.15 | 0.81 | 1.05 | 0.08 |
| M87 SMA 40 | 47.19 | 67.98 | 1.34 | 0.19 |
| M87 DIBMA | 27.78 | 4.44 | 0.37 | 0.19 |
| M87 Sulfo-DIBMA | 30.74 | 2.22 | 0.34 | 0.12 |
| M87 Sulfo-SMA | 32.47 | 1.77 | 0.30 | 0.05 |
| M87 SMA BZ25 | 354.11 | 877.45 | 1.37 | 0.26 |
| M87 SMA BZ30 | 9.76 | 1.04 | 0.85 | 0.07 |
| M87 SMA BZ35 | 10.97 | 0.50 | 0.63 | 0.04 |
| M87 SMA BZ40 | 12.13 | 3.13 | 0.75 | 0.12 |
| M87 AASTY 6-45 | 12.97 | 7.95 | 1.10 | 0.08 |
| M87 AASTY 11-45 | 10.07 | 2.43 | 0.91 | 0.05 |
| M87 AASTY 6-50 | 11.23 | 2.50 | 0.68 | 0.15 |
| M87 AASTY 11-50 | 9.20 | 1.02 | 0.74 | 0.06 |
| M87 AASTY 6-55 | 10.82 | 6.70 | 0.78 | 0.35 |
| M87 AASTY 11-50 | 13.04 | 0.44 | 0.38 | 0.03 |

**Figure S7. Polymer screen data for *S. muelleri* BiPGT1. (A)** Radius and polydispersity data for polymers in the following order: SMA100, SMA200, SMA300, DIBMA, Sulfo-DIBMA, Sulfo-SMA, SMA BZ25, SMA BZ30, SMA BZ35, SMA BZ40, AASTY 6-45, AASTY 11-45, AASTY 6-50, AASTY 11-50, AASTY 6-55, AASTY 11-55. Due to low protein yield SDS-PAGE revealed faint bands precluding reliable interpretation, thus DLS data was used for designating the optimal polymer for BiPGT2.

**A**

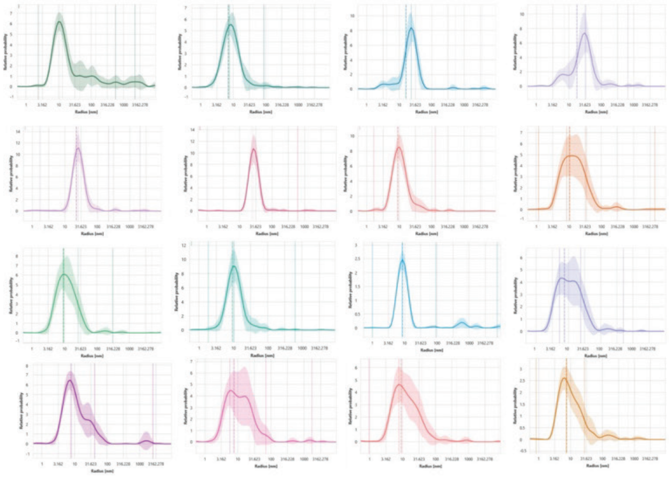

| Annotations | Cumulant Fit | | | |
|---|---|---|---|---|
| | Cumulant Radius (nm) | | Cumulant PDI | |
| Sample ID | ø | σ | ø | σ |
| PglI SMA25 | 13.13 | 1.35 | 0.63 | 0.12 |
| PglI SMA30 | 7.19 | 0.08 | 0.37 | 0.02 |
| PglI SMA40 | 16.31 | 0.26 | 0.46 | 0.02 |
| PglI DIBMA | 17.78 | 0.44 | 0.57 | 0.03 |
| PglI Sulfo-DIBMA | 23.04 | 0.55 | 0.28 | 0.02 |
| PglI Sulfo-SMA | 24.81 | 0.56 | 0.24 | 0.02 |
| PglI SMALP BZ25 | 9.07 | 0.09 | 0.30 | 0.01 |
| PglI SMALP BZ30 | 10.69 | 0.34 | 0.49 | 0.05 |
| PglI SMALP BZ35 | 9.48 | 0.12 | 0.40 | 0.02 |
| PglI SMALP BZ40 | 9.45 | 0.13 | 0.31 | 0.01 |
| PglI AASTY 6-45 | 8.79 | 0.13 | 0.33 | 0.03 |
| PglI AASTY 11-45 | 7.93 | 0.18 | 0.52 | 0.02 |
| PglI AASTY 6-50 | 8.18 | 0.66 | 0.48 | 0.11 |
| PglI AASTY 11-50 | 7.71 | 0.21 | 0.56 | 0.03 |
| PglI AASTY 6-55 | 9.21 | 0.19 | 0.53 | 0.03 |
| PglI AASTY 11-55 | 7.57 | 0.25 | 0.47 | 0.03 |

**B**

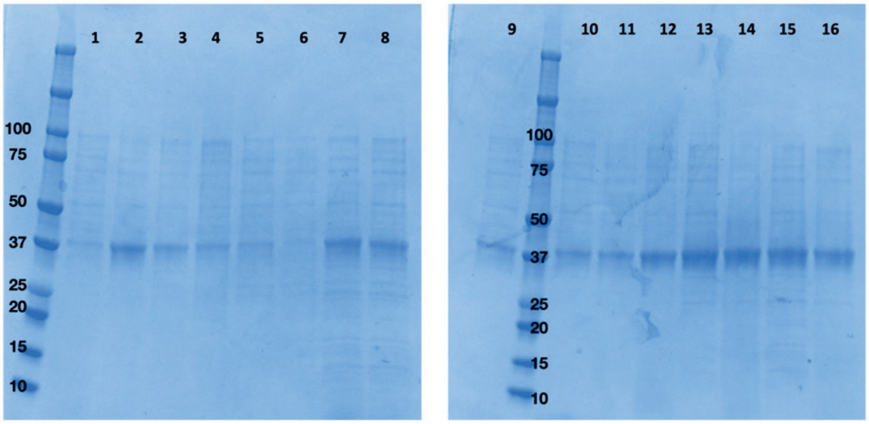


**Figure S8. Polymer screen data for *C. concisus* PglI. (A)** Radius and polydispersity data for polymers in the following order: SMA100, SMA200, SMA300, DIBMA, Sulfo-DIBMA, Sulfo-SMA, SMA BZ25, SMA BZ30, SMA BZ35, SMA BZ40, AASTY 6-45, AASTY 11-45, AASTY 6-50, AASTY 11-50, AASTY 6-55, AASTY 11-55. **(B)** SDS-PAGE of purification of PglI using the polymer screen. Lane numbering is the same as Fig. S3.

**A**

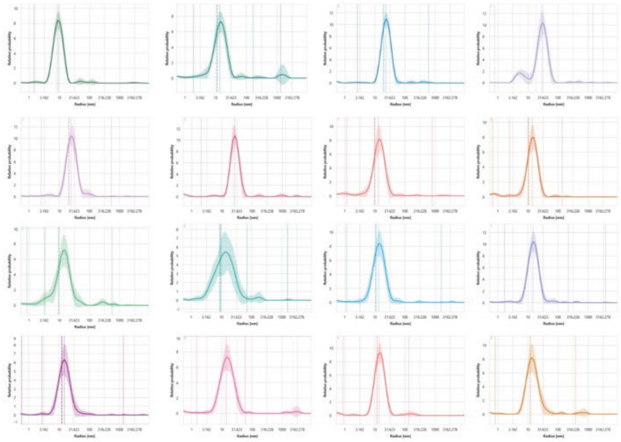

| Annotations | Cumulant Fit | | | |
|---|---|---|---|---|
| | Cumulant Radius (nm) | | Cumulant PDI | |
| Sample ID | ø | σ | ø | σ |
| GT1 SMA25 | 11.32 | 1.83 | 0.49 | 0.16 |
| GT1 SMA30 | 19.78 | 0.35 | 0.32 | 0.02 |
| GT1 SMA40 | 18.75 | 0.39 | 0.53 | 0.02 |
| GT1 DIBMA | 21.08 | 0.33 | 0.33 | 0.02 |
| GT1 Sulfo-DIBMA | 22.57 | 0.68 | 0.34 | 0.03 |
| GT1 Sulfo-SMA | 10.03 | 0.38 | 0.48 | 0.03 |
| GT1 SMALP BZ25 | 10.41 | 0.24 | 0.48 | 0.03 |
| GT1 SMALP BZ30 | 9.50 | 0.26 | 0.56 | 0.02 |
| GT1 SMALP BZ35 | 9.40 | 0.11 | 0.50 | 0.02 |
| GT1 SMALP BZ40 | 11.01 | 0.24 | 0.36 | 0.02 |
| GT1 AASTY 6-45 | 12.72 | 0.17 | 0.30 | 0.01 |
| GT1 AASTY 11-45 | 12.67 | 0.24 | 0.34 | 0.03 |
| GT1 AASTY 6-50 | 12.03 | 0.80 | 0.44 | 0.07 |
| GT1 AASTY 11-50 | 11.98 | 0.26 | 0.37 | 0.03 |
| GT1 AASTY 6-55 | 12.08 | 0.60 | 0.35 | 0.05 |
| GT1 AASTY 11-55 | 11.32 | 1.83 | 0.49 | 0.16 |

**B**

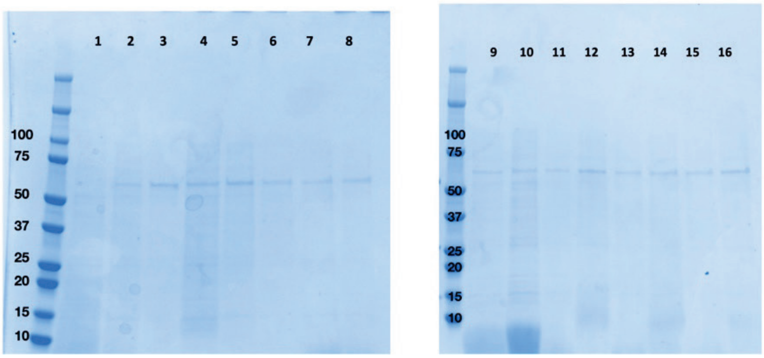


**Figure S9. Polymer screen data for *Myxococcales bacterium* GT1. (A)** Radius and polydispersity data for polymers in the following order: SMA100, SMA200, SMA300, DIBMA, Sulfo-DIBMA, Sulfo-SMA, SMA BZ25, SMA BZ30, SMA BZ35, SMA BZ40, AASTY 6-45, AASTY 11-45, AASTY 6-50, AASTY 11-50, AASTY 6-55, AASTY 11-55. **(B)** SDS-PAGE of purification of GT1 using the polymer screen. Lane numbering is the same as Fig. S3. Due to the similarity in molecular weight with Gro-EL, the difference in radius was utilized to show that the signal corresponds to GT1.

**A**

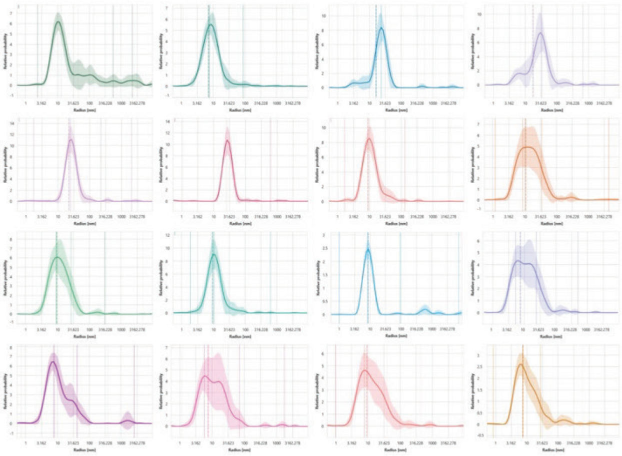

| Annotations | Cumulant Fit | | | |
|---|---|---|---|---|
| | Cumulant Radius (nm) | | Cumulant PDI | |
| Sample ID | ø | σ | ø | σ |
| WecP SMA25 | 8.08 | 0.12 | 0.34 | 0.01 |
| WecP SMA30 | 9.35 | 0.21 | 0.30 | 0.03 |
| WecP SMA40 | 20.36 | 0.99 | 0.32 | 0.07 |
| WecP DIBMA | 16.76 | 0.28 | 0.50 | 0.03 |
| WecP Sulfo-DIBMA | 22.80 | 1.20 | 0.26 | 0.10 |
| WecP Sulfo-SMA | 23.71 | 0.57 | 0.30 | 0.03 |
| WecP SMALP BZ25 | 9.82 | 0.13 | 0.43 | 0.02 |
| WecP SMALP BZ30 | 10.42 | 0.13 | 0.42 | 0.01 |
| WecP SMALP BZ35 | 10.18 | 0.25 | 0.40 | 0.03 |
| WecP SMALP BZ40 | 9.17 | 0.09 | 0.31 | 0.03 |
| WecP AASTY 6-45 | 11.02 | 0.40 | 0.30 | 0.08 |
| WecP AASTY 11-45 | 9.12 | 0.11 | 0.35 | 0.03 |
| WecP AASTY 6-50 | 9.98 | 0.43 | 0.29 | 0.07 |
| WecP AASTY 11-50 | 9.30 | 0.12 | 0.40 | 0.01 |
| WecP AASTY 6-55 | 8.70 | 0.28 | 0.34 | 0.04 |
| WecP AASTY 11-55 | 8.42 | 0.08 | 0.28 | 0.01 |

**B**

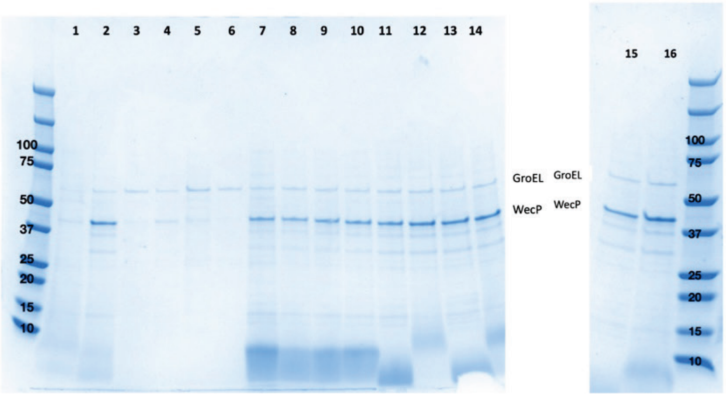


**Figure S10. Polymer screen data for *A. hydrophila* WecP. (A)** Radius and polydispersity data for polymers in the following order: SMA100, SMA200, SMA300, DIBMA, Sulfo-DIBMA, Sulfo-SMA, SMA BZ25, SMA BZ30, SMA BZ35, SMA BZ40, AASTY 6-45, AASTY 11-45, AASTY 6-50, AASTY 11-50, AASTY 6-55, AASTY 11-55. **(B)** SDS-PAGE of purification of WecP using the polymer screen. Lane numbering is the same as Fig. S3.

**A**

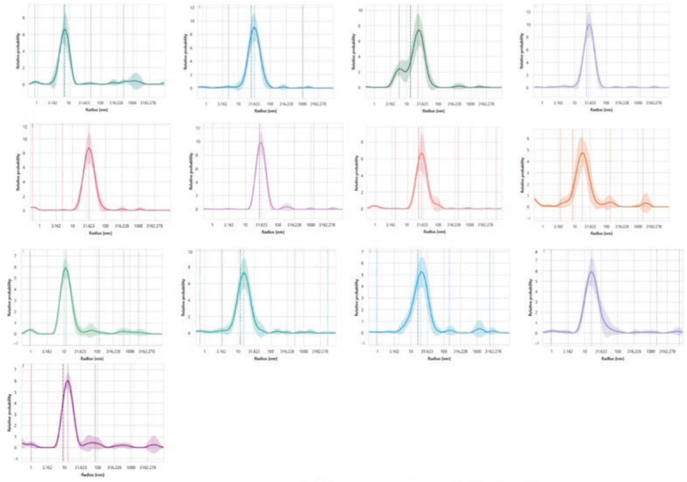

| Annotations | Cumulant Fit | | | |
|---|---|---|---|---|
| | Cumulant Radius (nm) | | Cumulant PDI | |
| Sample ID | ø | σ | ø | σ |
| Tt WbaP SMA25 | 7.19 | 1.84 | 0.44 | 0.28 |
| Tt WbaP SMA30 | 13.56 | 0.35 | 0.53 | 0.01 |
| Tt WbaP SMA40 | 25.63 | 0.85 | 0.36 | 0.03 |
| Tt WbaP DIBMA | 24.13 | 0.51 | 0.31 | 0.03 |
| Tt WbaP Sulfo-DIBMA | 27.00 | 0.56 | 0.33 | 0.02 |
| Tt WbaP Sulfo-SMA | 27.44 | 0.69 | 0.24 | 0.04 |
| Tt WbaP SMALP BZ25 | 22.89 | 0.86 | 0.43 | 0.04 |
| Tt WbaP SMALP BZ30 | 8.86 | 2.11 | 1.05 | 0.08 |
| Tt WbaP AASTY 6-45 | 10.78 | 0.39 | 0.47 | 0.07 |
| Tt WbaP AASTY 11-45 | 16.27 | 1.06 | 0.45 | 0.11 |
| Tt WbaP AASTY 6-50 | 11.99 | 0.27 | 0.46 | 0.02 |
| Tt WbaP AASTY 11-50 | 14.35 | 1.11 | 0.45 | 0.10 |
| Tt WbaP AASTY 11-55 | 9.37 | 3.72 | 0.84 | 0.30 |

**B**

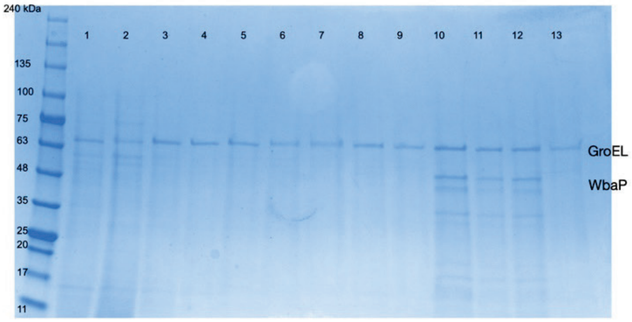


**Figure S11. Polymer screen data for *T. thermophilus* WbaP. (A)** Radius and polydispersity data for polymers in the following order: SMA100, SMA200, SMA300, DIBMA, Sulfo-DIBMA, Sulfo-SMA, SMA BZ25, SMA BZ30, AASTY 6-45, AASTY 11-45, AASTY 6-50, AASTY 11-50, AASTY 11-55. **(B)** SDS-PAGE of purification of WbaP using the polymer screen. Lanes numbers correspond to samples: **1**: SMA100, **2**: SMA200, **3:** SMA300, **4**: DIBMA, **5**: Sulfo-DIBMA, **6**: Sulfo-SMA, **7**: SMA BZ25, **8**: SMA BZ30, **9**: AASTY 6-45, **10**: AASTY 11-45, **11:** AASTY 6-50, **12**: AASTY 11-50, **13**: AASTY 11-55.

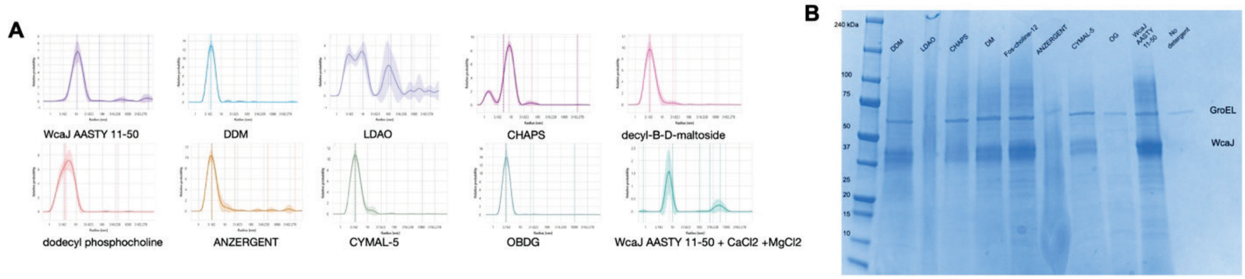


**Figure S12. Detergent exchange screen data for *E. coli* WcaJ. (A)** DLS data demonstrates destabilization of lipid nanoparticle and exchange into detergent micelles by a change in radius from ~10 nm to ~4 nm. **(B)** SDS-PAGE confirms successful exchange of WcaJ from lipid nanoparticle into detergent micelles.

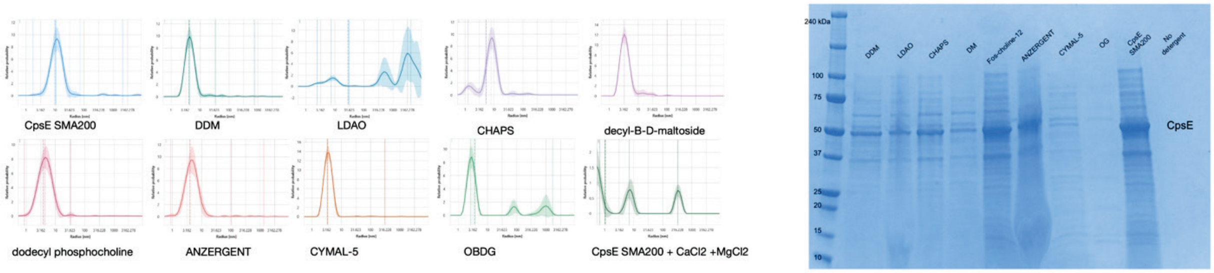


**Figure S13. Detergent exchange screen data for *S. pneumoniae* CpsE. (A)** DLS data demonstrates destabilization of lipid nanoparticle and exchange into detergent micelles by a change in radius from ~10 nm to ~4 nm. **(B)** SDS-PAGE confirms successful exchange of CpsE from lipid nanoparticle into detergent micelles.

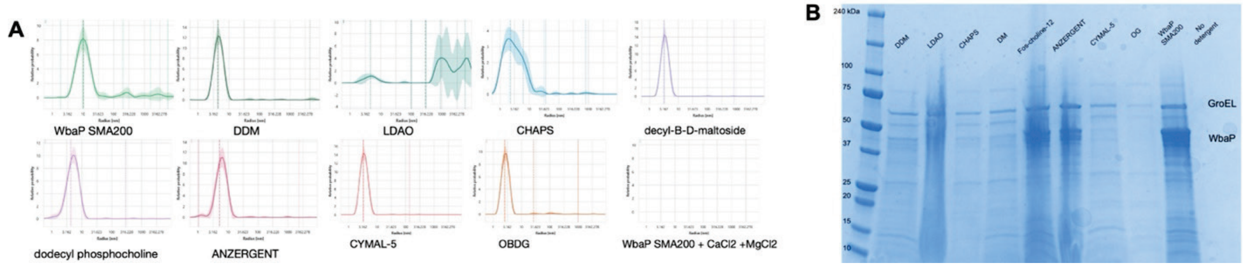


**Figure S14. Detergent exchange screen data for *S. enterica* WbaP. (A)** DLS data demonstrates destabilization of lipid nanoparticle and exchange into detergent micelles by a change in radius from ~10 nm to ~4 nm. **(B)** SDS-PAGE confirms successful exchange of WbaP from lipid nanoparticle into detergent micelles.

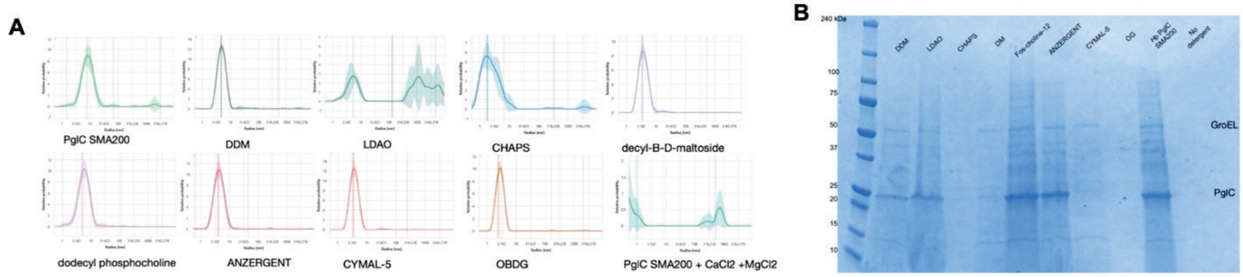


**Figure S15. Detergent exchange screen data for *H. pullorum* PglC. (A)** DLS data demonstrates destabilization of lipid nanoparticle and exchange into detergent micelles by a change in radius from ~10 nm to ~4 nm. **(B)** SDS-PAGE confirms successful exchange of PglC from lipid nanoparticle into detergent micelles.

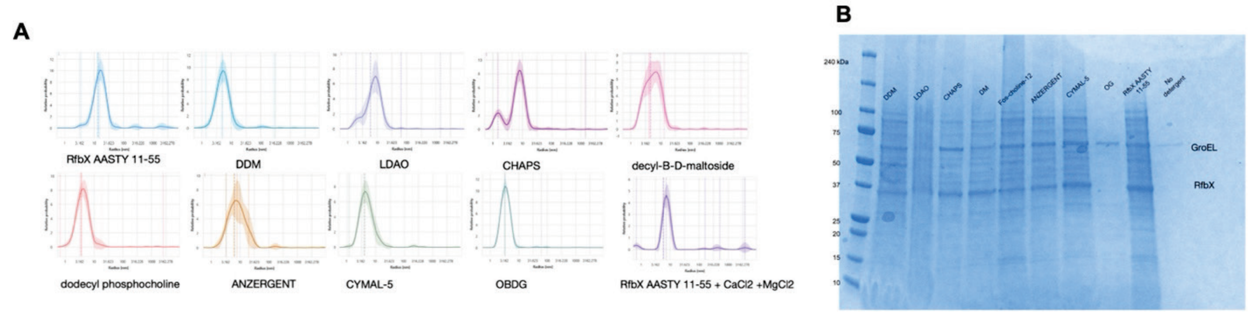


**Figure S16. Detergent exchange screen data for *S. enterica* RfbX. (A)** DLS data demonstrates destabilization of lipid nanoparticle and exchange into detergent micelles by a change in radius from ~10 nm to ~4 nm. **(B)** SDS-PAGE confirms successful exchange of RfbX from lipid nanoparticle into detergent micelles.

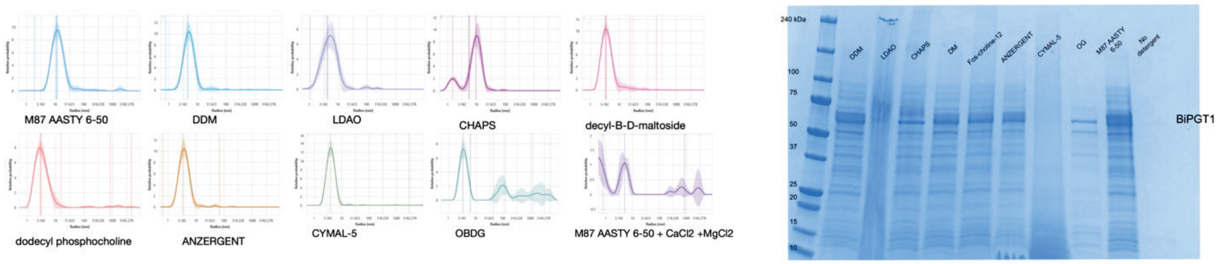


**Figure S17. Detergent exchange screen data for *S. muelleri* BiPGT1. (A)** DLS data demonstrates destabilization of lipid nanoparticle and exchange into detergent micelles by a change in radius from ~10 nm to ~4 nm. **(B)** SDS-PAGE confirms successful exchange of BiPGT1 from lipid nanoparticle into detergent micelles.

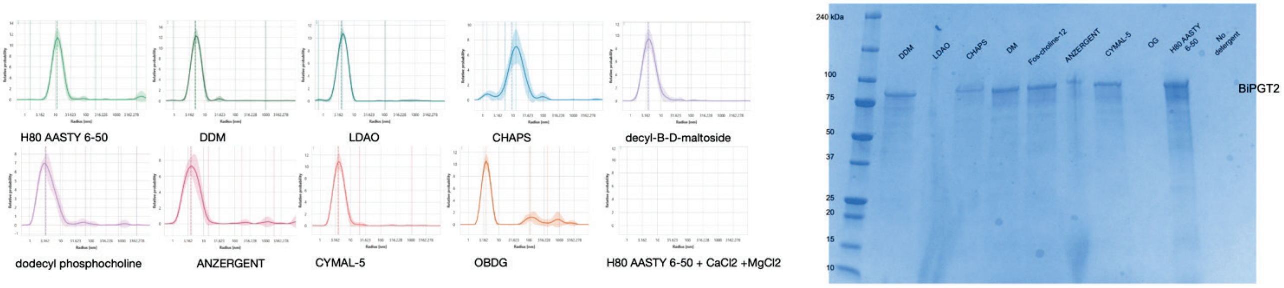


**Figure S18. Detergent exchange screen data for *Limnohabitans sp. Jir61* BiPGT2. (A)** DLS data demonstrates destabilization of lipid nanoparticle and exchange into detergent micelles by a change in radius from ~10 nm to ~4 nm. **(B)** SDS-PAGE confirms successful exchange of BiPGT2 from lipid nanoparticle into detergent micelles.

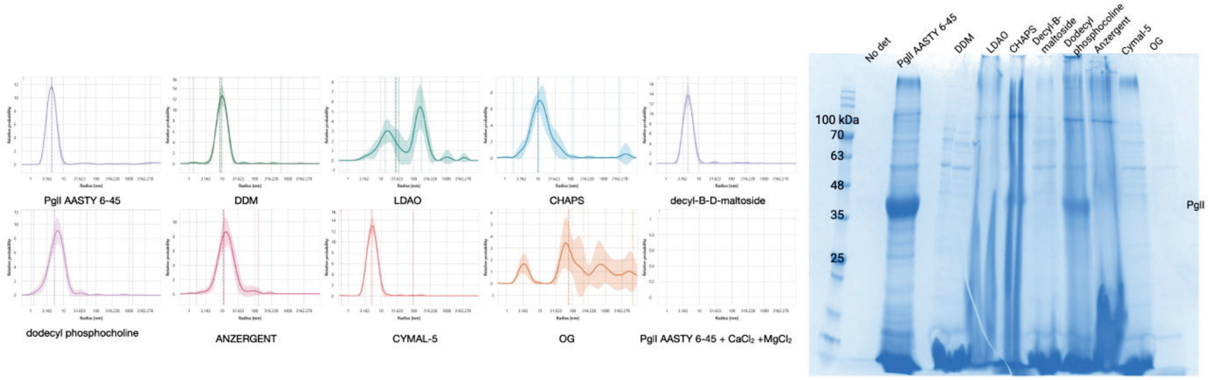


**Figure S19. Detergent exchange screen data for *C. concisus* PglI. (A)** DLS data demonstrates destabilization of lipid nanoparticle and exchange into detergent micelles by a change in radius from ~10 nm to ~4 nm. **(B)** SDS-PAGE confirms successful exchange of PglI from lipid nanoparticle into detergent micelles.

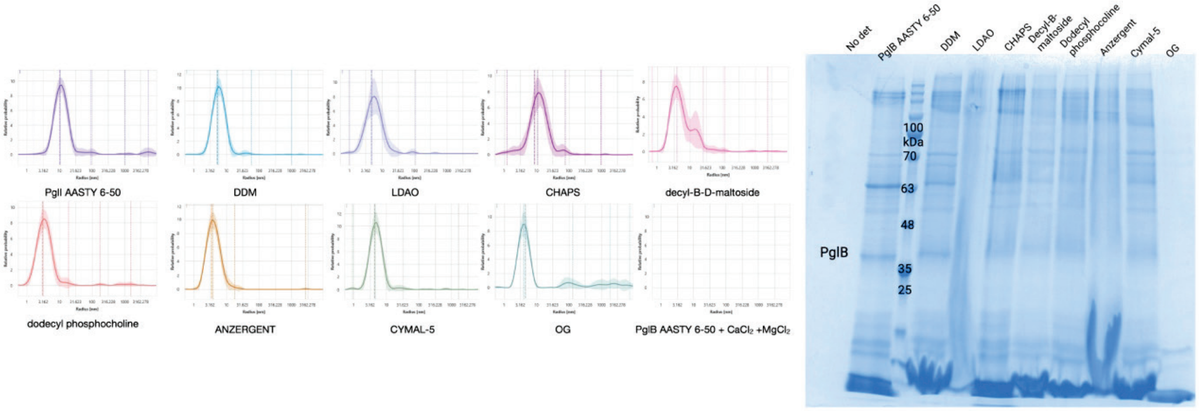


**Figure S20. Detergent exchange screen data for *N. gonorrhoeae* PglB. (A)** DLS data demonstrates destabilization of lipid nanoparticle and exchange into detergent micelles by a change in radius from ~10 nm to ~4 nm. **(B)** SDS-PAGE confirms successful exchange of PglB from lipid nanoparticle into detergent micelles.

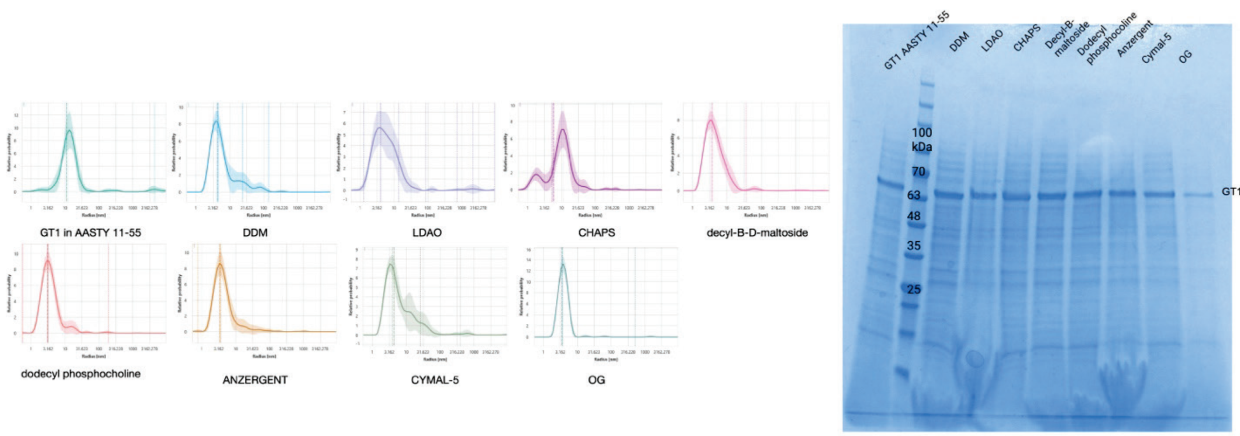


**Figure S21. Detergent exchange screen data for *Myxococcales bacterium* GT1. (A)** DLS data demonstrates destabilization of lipid nanoparticle and exchange into detergent micelles by a change in radius from ~10 nm to ~4 nm. **(B)** SDS-PAGE confirms successful exchange of GT1 from lipid nanoparticle into detergent micelles.

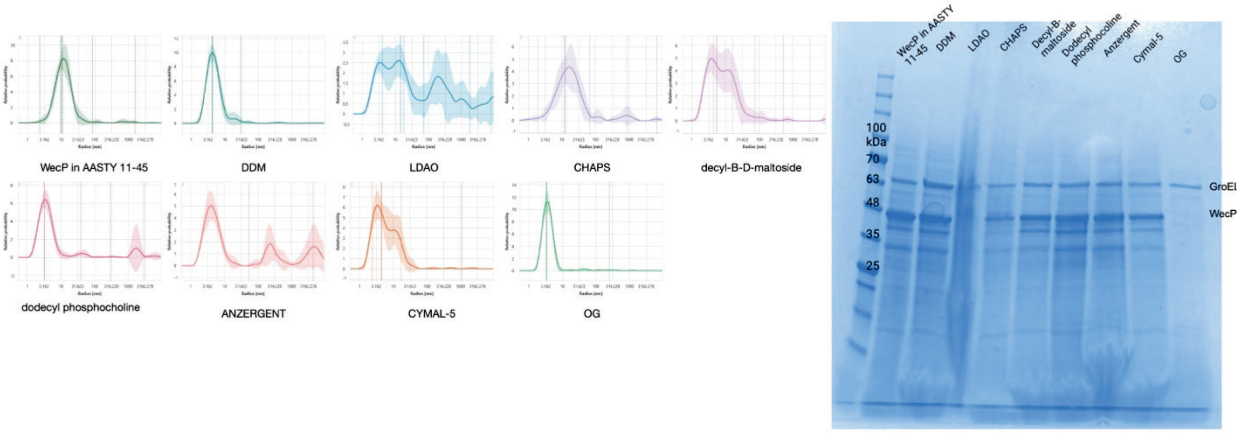


**Figure S22. Detergent exchange screen data for *A. hydrophila* WecP. (A)** DLS data demonstrates destabilization of lipid nanoparticle and exchange into detergent micelles by a change in radius from ~10 nm to ~4 nm. **(B)** SDS-PAGE confirms successful exchange of WecP from lipid nanoparticle into detergent micelles.

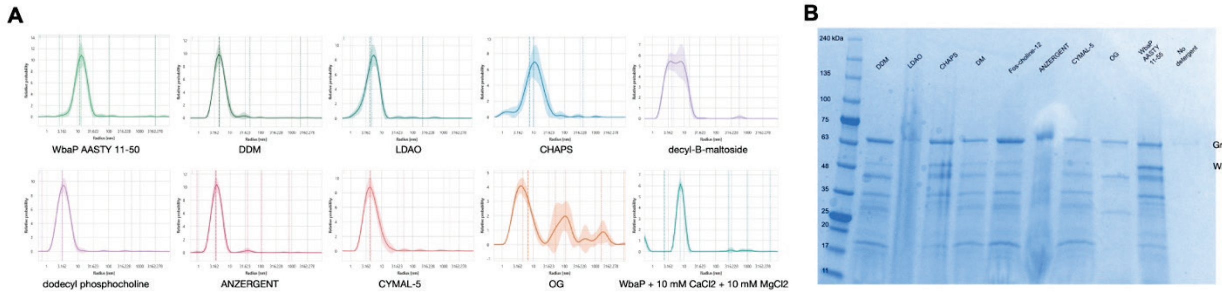


**Figure S23. Detergent exchange screen data for *T. thermophilus* WbaP. (A)** DLS data demonstrates destabilization of lipid nanoparticle and exchange into detergent micelles by a change in radius from ~10 nm to ~4 nm. **(B)** SDS-PAGE confirms successful exchange of WbaP from lipid nanoparticle into detergent micelles.

### Table S1. Primers for cloning plasmids.

| Primer name | Primer sequence 5' -> 3' |
|---|---|
| **SST Vector Fwd** | CTCGAGTGAGATCCGGCTGC |
| **SST Vector Rev** | TTGCATTGGATTGGAAGTACAGGTT |
| ***T.t.* WbaP Fwd** | GTACTTCCAATCCAATGCAGGCACGGCCCCTC |
| ***T.t.* WbaP Rev** | GCAGCCGGATCTCACTCGAGTTAGTATGCGCCTTTTCTAGTCAGGACCAC |
| **BiPGT1 Fwd** | GGGTACCGAGAACCTGTACTTCCAATCCAATGCAATGAAGTTTTCGGTACTCATG |
| **BiPGT1 Rev** | GCAGCCGGATCTCACTCGAGTTAACGAGCCCCAAAACC |
| **BiPGT2 Fwd** | GTACTTCCAATCCAATGCAATGGTTCCTCATGTCATGATTG |
| **BiPGT2 Rev** | GCAGCCGGATCTCACTCGAGTCATCTGGCCCCAAATCC |
| ***C.j.* PglI Fwd** | TGTACTTCCAATCCAATGCAATGCCAAAGCTTAGTGTCATCGT |
| ***C.j.* PglI Rev** | GCAGCCGGATCTCACTCGAGTCAGTTTTTGCAC |
| ***C.c.* 13826 PglI Fwd** | AGATCTGGGTACCGAGAACCTGTACTTCCAATCCAATGCAATGATGTCTGAGCCGCTGAT |
| ***C.c.* 13826 PglI Rev** | CTTCCTTTCGGGCTTTGTTAGCAGCCGGATCTCACTCGAGTCATACCCCGAAACGCTGTTTTATTTTAACG |
| **GT1 Fwd** | TGTACTTCCAATCCAATGCAATGAGCATCGGTATTGACATCCTGA |
| **GT2 Rev** | GCAGCCGGATCTCACTCGAGTTAGTGCGCACC |
